# ABCA1 and apoA-I dependent 12-hydroxyeicosatetraenoic acid efflux regulates macrophage inflammatory signaling

**DOI:** 10.1101/2024.07.11.603001

**Authors:** Brian A. Harsch, Kamil Borkowski, Rachel E. Walker, Theresa L. Pedersen, John W. Newman, Gregory C. Shearer

## Abstract

Aberrant high-density lipoprotein (HDL) function is implicated in inflammation-associated pathologies. While HDL ABCA1-mediated reverse cholesterol and phospholipid transport are well described, the movement of pro-/anti-inflammatory lipids has not been explored. HDL phospholipids are the largest reservoir of circulating arachidonic acid-derived oxylipins. Endotoxin-stimulation activates inflammatory cells leading to hydroxyeicosatetraenoic acid (HETE) production, oxylipins which are involved in inflammatory response coordination. Active signaling in the non-esterified (NE) pool is terminated by sequestration of HETEs as esterified (Es) forms and degradation. We speculate that an ABCA1-apoA-I-dependent efflux of HETEs from stimulated cells could regulate intracellular HETE availability. Here we test this hypothesis both in vitro and in vivo. In endotoxin-stimulated RAW-264.7 macrophages preloaded with d8-arachidonic acid we use compartmental tracer modeling to characterize the formation of HETEs, and their efflux into HDL. We found that in response to endotoxin: I) Cellular NE 12-HETE is positively associated with MCP-1 secretion (p<0.001); II) HETE transfer from NE to Es pools is ABCA1-depedent (p<0.001); III) Cellular Es HETEs are transported into media when both apoA-I and ABCA1 are present (p<0.001); IV) The stimulated efflux of HETEs >> arachidonate (p<0.001). Finally, in endotoxin challenged humans (n=17), we demonstrate that intravenous lipopolysaccharide (0.6 ng/kg body weight) resulted in accumulation of 12-HETE in HDL over a 168-hour follow-up. Therefore, HDL can suppress inflammatory responses in macrophages by regulating intracellular HETE content in an apoA-I/ABCA1 dependent manner. The described mechanism may apply to other oxylipins and explain anti-inflammatory properties of HDL. This newly defined HDL property opens new doors for the study of lipoprotein interactions in metabolic diseases.

## Introduction

HDL are known for their role in reverse cholesterol transport and circulating HDL cholesterol is inversely associated with hazards for cardiovascular, metabolic, and cognitive diseases and more^1–5^. In the prior decade, intense clinical trial activity tested multiple pharmaceuticals aimed at raising HDL-cholesterol (e.g. extended-release niacin^6^, torcetrapib^7^), however no therapeutic benefits emerged highlighting the need for a better understanding of HDL function^8^. In response, we considered the implications of HDL as a transporter of oxylipins since HDL constitute the largest pool of circulating oxylipins derived from arachidonic, eicosapentaenoic, and docosahexaenoic acids^9^. Among many activities, oxylipins are regulators of both acute and low-grade systemic inflammation and energy metabolism^10^. Our previous efforts have demonstrated that HDL oxylipin content is altered by metabolic disorders such as metabolic syndrome^11^, alterations in dietary PUFA content^12^, and pharmacological interventions^11–13^. While the origins of HDL-oxylipins are unknown, their bioactivities and presence in HDL suggest an unaccounted-for element of HDL biology.

Oxylipins are broad suite of regulatory metabolites generated by the action of lipoxygenase (LOX), cyclooxygenase (COX), and cytochrome 450 (CYP), or autooxidation^14^. Canonically, oxylipins are formed intracellularly, following activation of cPLA_2_ and the release of PUFAs from the *sn-2* position on phospholipids^15^. While true for COX and CYP metabolism, LOXs are also surface acting enzymes that can directly oxidize membrane PUFAs, which can then be released by cPLA_2_ ^16–18^. Non-esterified (NE, i.e., “free”) oxylipins are ligands for GPCRs, PPAR family receptors, and more, with specificity depending on the form and location of oxygen incorporation into the PUFA molecule. For instance, in macrophages, the response to lipopolysaccharide (LPS) inflammatory challenge to is mediated by intracellular 12-HETE which activates protein kinase c (PKC), leading to subsequent MCP-1 production^19^. The phospholipid *sn-2* position is also the site for re-esterification of oxylipins, which facilitates signal termination^19^. It is thought oxylipins participate in recycling between the non-esterified and esterified pools by participation in the Lands cycle^20,21^. Notably, hydroxy and epoxy fatty acids are substrates for long chain fatty acid CoA ligase 1 (ACSL1) and ACSL4^22^. Interestingly, the macrophage inflammatory response to high fat feeding is dependent on ASCL1 activating CD36-FABP4-p38-PPARd^23^. Therefore, recycling may be a necessary element of oxylipin activity, amplifying and extending the presence NE-oxylipins in the cytosol. However, in conditions where oxylipin accumulation in membrane phospholipids is excessive recycling would delay signal termination.

As a component of membrane phospholipids, oxylipins represent a point of overlap with HDL biology. While cholesterol is the most studied lipid transferred by the ABCA1/apoA-I exchange mechanisms, phospholipids are also essential components of lipid exchange^24^. Hence, if phospholipids containing oxylipins are exchanged, it would represent a loss of esterified oxylipins from the recycling oxylipin pool, and consequently indirectly reduce intracellular NE-oxylipins. Such a dynamic would provide a molecular mechanism to explain the link between HDL biology and its inverse association with chronic inflammatory conditions.

Here we test whether the appearance of oxylipins in nascent HDL is ABCA1- or apoA-I-dependent and affect intracellular NE-oxylipin content and downstream inflammatory outcomes. We use a simple cell model of inflammation, the LPS-challenged RAW-264.7 mouse macrophage in which we trace deuterium-labeled arachidonic acid (d8-AA) from the cell esterified fraction through the elements of our construct: the non-esterified (NE) arachidonic acid (AA) fraction, lipoxygenase-mediated conversion to intracellular NE-HETEs, re-esterification of HETEs into the cell-esterified (Es) fraction, and transfer of Es-HETEs from the cell to the media. We do this in a 2×2 factorial manner with conditions lacking both apoA-I and ABCA1, conditions lacking either ABCA1 or apoA-I, and in the full system expressing ABCA1 and exposed to apoA-I. Analytically, we use compartmental modeling to measure the characteristic transfer of three specific d8-AA-oxylipins derived from LOXs: d8-5-, d8-12-, and d8-15-HETE produced by 5-LOX and 12/15-LOX respectively. Using this approach: ***a)*** we observe movement of label through each measured pool; ***b)*** we compare essential metrics of oxylipin exchange as fractional transfer rates (FTRs), area-under the curve (AUC), and intercompartmental flux (*Φ*); and ***c)*** we relate them to MCP-1, an inflammatory cytokine whose expression in macrophages is elicited by NE-12-HETEs. Finally, to evaluate the relevance of our findings at the organismal level, we support the intracellular model with HDL-HETE responses in endotoxin-challenged humans.

## Results

### I: Model system suitability

To support the modeling effort, we first evaluated the quality of the cell systems. The effective comparison of model features requires successful silencing of ABCA1 and sustainable cell viability in minimal conditions over a time period sufficient for experimental modeling. Effective ABCA1 protein and mRNA suppression, along with minimal effects of the endotoxin challenge on cell count, cell AA content, and cell palmitic acid content (a saturated fatty acid unrelated to HETE signaling) are demonstrated in **Supplemental Figure 1**.

### II: apoA-I/ABCA1 mediates selective HETE removal by membrane efflux to HDL

We next examined the overall effect of apoA-I and ABCA1 on the time-dependent changes of AA and HETE abundance in the four main pools examined: the NE and Es pools of the cell pellets (i.e., NE_cell_ and Es_cell_) and media (i.e., NE_media_ and Es_media_; **Supplemental Figure 2**). The presence of ABCA1 subtly increases the AA Es_cell_ pool, but the pool size itself is stable with time (**SFig2a**). In contrast, the d8-AA NE_cell_ pool is unaffected by either apoA-I or ABCA1 and is gradually depleted by ∼15% over the 3h experimental time course (**SFig2b**). In the extracellular pools, the presence of apoA-I enhances the ABCA1-dependent increase in AA abundance in the Es_media_ pool and in the NE_media_ pool (**SFig2c-d**). The HETEs generally tracked each other but showed distinct behavior compared to AA (**SFig2e-p**). In the intracellular pools, the accumulation of Es_cell_-HETE was amplified in the absence of apoA-I, while the accumulation of intracellular NE_cell_ HETEs in the absence of ABCA1 was rapid. In the extracellular pool, rapid accumulation of Es_media_-HETEs only occurred in the presence of both apoA-I and ABCA1, consistent with ABCA1-dependent transfer of phospholipids containing Es_media_-HETEs to facilitate the formation of nascent apoA-I HDL. While there was relatively little accumulation of NE_media_ HETE under any condition, it was most rapid in the absence of ABCA1, and the presence of apoA-I partially limited it.

### III: Intracellular NE-12-HETE mediates endotoxin-stimulated MCP-1 secretion

Using structural equation modeling mediation analysis, we confirm that HDL can mediate inflammatory outcomes by regulating intracellular NE-12-HETE concentrations (**Figure 1**). Our model, expressed in **Fig1a,** provides a statistical test to whether the effect of apoA-I and ABCA1 to reduce media MCP-1 accumulation occurs directly, irrespective of NE_cell_-12-HETE, or indirectly by modulating intracellular NE-12-HETE concentrations. The prerequisites for a mediation analysis using this model were met as: 1) a pre-existing causal mechanism for MCP-1 expression to be regulated by NE-12-HETE^19,25^; 2) the treatment condition predicts MCP-1 outcomes (i.e., apoA-I and ABCA1 suppress MCP-1 appearance in the media, **Fig1b**); 3) the treatment conditions predict the proposed mediator (i.e., apoA-I and ABCA1 reduce NE-12-HETE, **Fig1c**); and 4) the response and mediator levels are associated (i.e., MCP1 levels are positively correlated with d8 12-HETE, **Fig1d**). Therefore, the strengths of the direct and indirect association could be calculated relative to the cumulative association (**Fig1e**) and are expressed as a percent: the effect of ABCA1 to suppress MCP-1 accumulation was large and almost completely (>90%) explained by its effect on intracellular NE-12-HETE; i.e. the indirect pathway **(Fig1f)**. While apoA-I had a smaller, mostly direct association independent of intracellular NE-12-HETE (Fig1e), in combination with ABCA1 its effect was dominated through its effect on NE_cell_-12-HETE. Hence, in macrophages responding to an endotoxin challenge, apoA-I and ABCA-1 suppress the rate of MCP-1 accumulation through their effect on intracellular NE-12-HETE.

**Figure 1:**
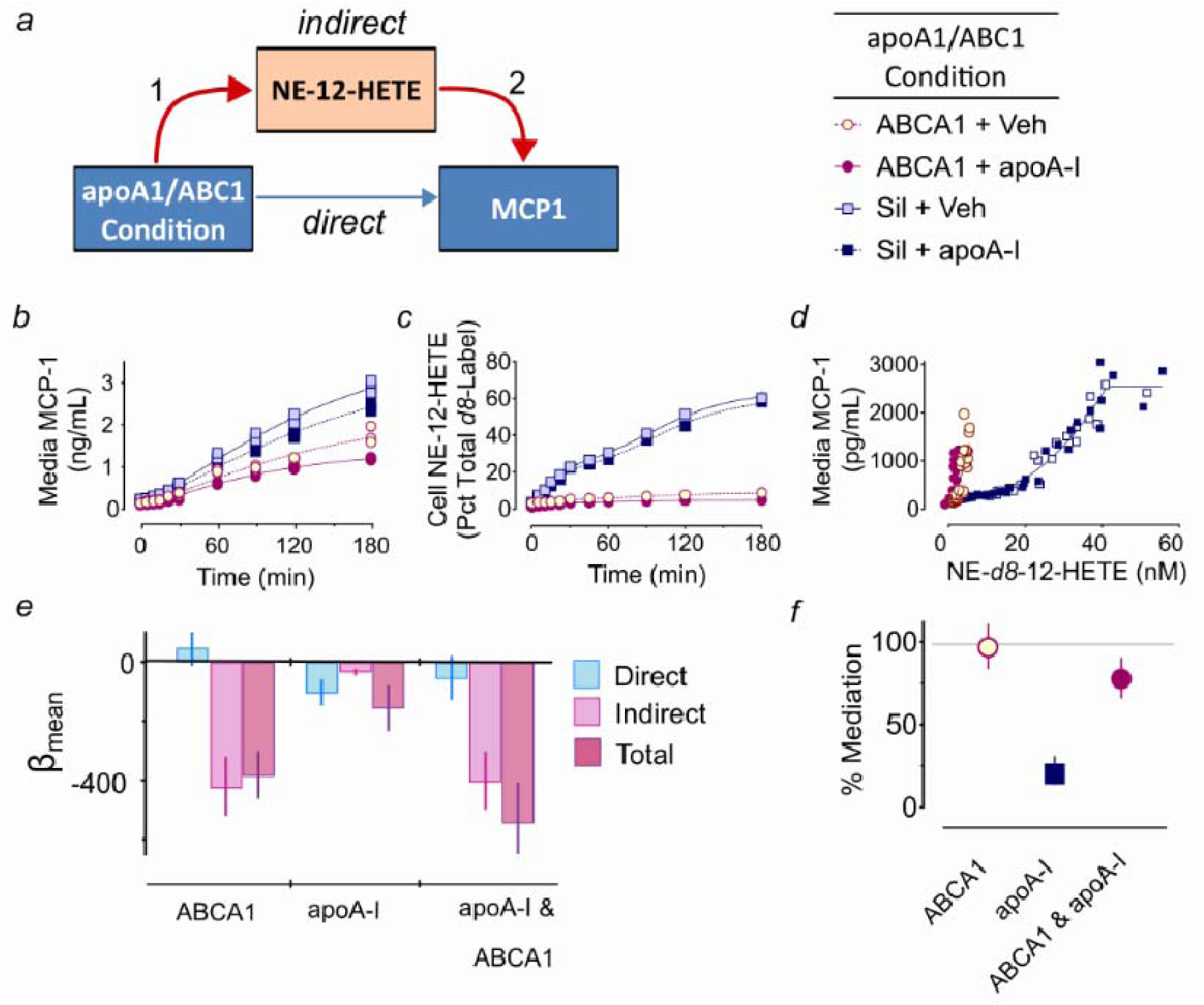
Intracellular NE-12-HETE mediates the association of condition with appearance of MCP-1 in the media. We used mediation analysis as a means to strengthen the causal inference – that apoA-I and ABCA1-dependent effects on MCP-1 appearance in media are explained by the effect of apoA-I and ABCA1 on intracellular 12-HETE concentrations. We based our analysis on the pathway construct **(A)** and confirmed the prerequisites of mediation: *1)* a significant association between apoA-I and ABCA1 status and media MCP-1 **(B)**; *2)* a significant association between apoA-I and ABCA1 status and intracellular NE-12-HETE **(C**, *same data as* ***SF1G*)**; and *3)* a significant association between intracellular NE-12-HETE and media MCP-1 **(D)**. We tested the contribution of the direct and indirect pathways as explanations of media MCP-1 using structural equations in which the total effect of treatment conditions are the sum of the direct contribution plus the indirect contribution. The direct effects (blue) are represented in comparison to the total contribution (purple) **(E)** and demonstrate that in both conditions having ABCA1 expressing status, there was very little direct contribution to the total association of condition with MCP-1. In contrast, the indirect pathway (pink) **(F)** provided a strong association with suppressing media MCP-1 appearance which accounted completely for the total effect **(G)** with veh/ABCA1 status, and almost completely with apoA-I/ABCA1 status. In contrast, in the apoA-I/Silenced condition, there was a small but significant suppressive contribution of apoA-I, but intracellular NE-12-HETE provided an explanation for only 20% of the total effect.

**Figure 2:**
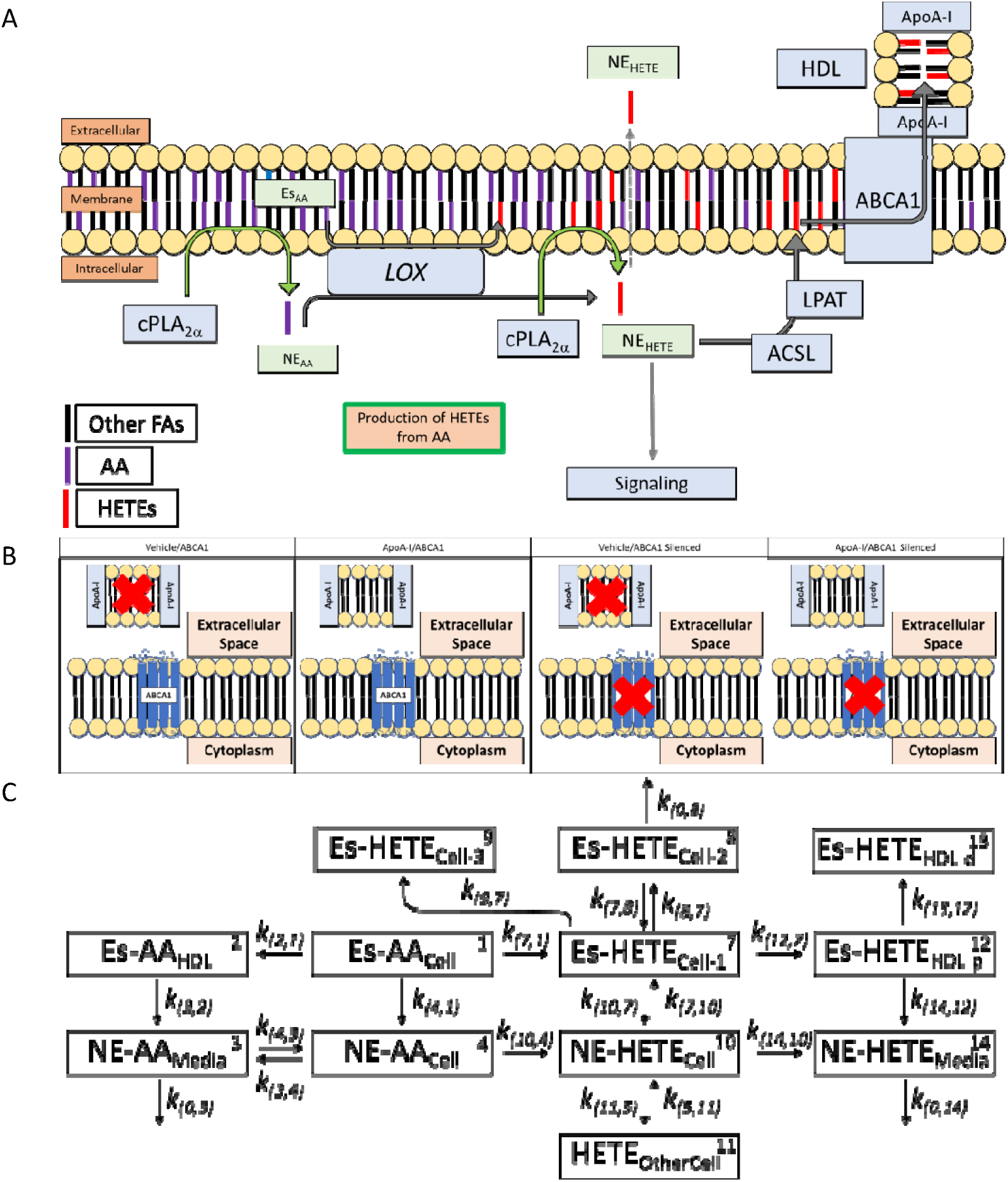
Conceptual framework and SAAM II model. **A)** Esterified 12-HETE is cleaved by cPLA_2_ to form 12-HETE. Non-esterified 12-HETE is active and can lead to increased MCP-1 production and is inactivated through re-esterification where it is effluxed to HDL for transport away from the cell. **B)** Cell experiment design. **C)** SAAM compartmental model for tracing d8-AA and d8-5-HETE, d8-12-HETE, and d8-15-HETE. The theoretical metabolite pools are numbered arbitrarily. By convention, kinetic constants described by k(to,from). Abbreviations: AA = arachidonic acid; Es = esterified; NE = nonesterified; HDL-p = proximal HDL pool; HDL-d = distal HDL pool;

### IV: Intercompartmental HETE transfer and consequential model features

#### Model fit and performance

The model construct in **F2c** traces label transfer between intracellular Es and NE pools, extracellular Es and NE (i.e., HDL and media) pools, along with other theoretical sub-compartments [e.g. intra and extracellular disposal, HDL-proximal (HDL-p) and -distal (HDL-d) pools] and is generally concordant with the absolute concentrations shown in **Supplemental Figure 2**. A single model including all pools and conditions allowed us to directly compare FTRs and maximize parsimony and model stability. Modeling the negative control condition alone [i.e., the Vehicle with ABCA1-silencing; (veh + Sil)] was not mathematically stable, even when sharing parameters from other conditions. The poor performance of this condition likely represents adaptive changes beyond the scope of our study and we do not emphasize the (veh + Sil) results or comparisons. We considered reports that HETE synthesis also occurs at multiple locations, classically in the cytosol [i.e., F2C, *k(10,4)*] or by direct oxygenation of PL-AA with phospholipid bilayers [i.e., F2C, *k(7,1)*]. We fit models detailing 5-HETE conversion in the membrane and in the cytosol. The models detailing cytosolic conversion not only lacked concordance with reported conversion locations ^26,27^, the transit and recycling values associated with 5-HETE trafficking was highly inconsistent with 12-HETE and 15-HETE; hence despite acceptable mathematical fit, we deemed the cytosolic appearance of 5-HETE lacked biological rigor. No models in which membrane conversion of 12-HETE or 15-HETE occurred with acceptable mathematical fit. In no case were FTRs associated with central to apoA-I/ABCA1-dependent HETE export substantively different. The final model fit with d8-AA, d8-5-HETE, d8-12-HETE, and d8-15-HETE was mathematically unique and sufficiently described the system without forfeiting information. The fit performance and model parameters are summarized in **Supplemental Table 1**. Generally, the Es_cell_ and NE_cell_ pools were well described; poor prediction was found in pools having small or no time-dependent changes in tracer abundance.

The fit for each pool is shown in **Figure 3**. The d8-AA is steadily depleted in both Es_cell_ (**F3a**) and NE_cell_ (**F3b**) pools under all conditions. Notably, in the presence of apoA-I and ABCA1, ∼8% of the total tracer appeared as AA in the extracellular Es_media_ pool (**F3c**) and ABCA1 alone allowed ∼0.5% into the extracellular NE_media_ pool (**F3d**). The behavior of the d8-HETEs were distinct from d8-AA, and subtly different from each other (**F3e-p**). All HETEs showed time-dependent accumulation in the Es_cell_-HETE pool that was greatest in the ABCA1 + veh condition; however, when apoA-I and ABCA1 were both present, tracer accumulation was nearly identical to Silenced cells (F3e,i,m). Silencing ABCA1 led to the rapid accumulation of label in the NE_cell_-HETE pool (F3f,j,n), while a saturating accumulation of HETEs in the Es_media_ pool occurred only in the presence of ABCA1 + apoA-I (F3g,k,o). As with the NE_cell_-HETE pool, the absence of ABCA1 and apoA-I resulted in slow accumulation of label in the NE_media_-HETE pool, which capped at ∼0.1% of the total tracer, or ∼10% of the intracellular NE-HETE levels (F3h,I,j). The fraction of NE-HETE transferred to the media was similar to that of NE-AA and may indicate facilitated diffusion.

**Figure 3:**
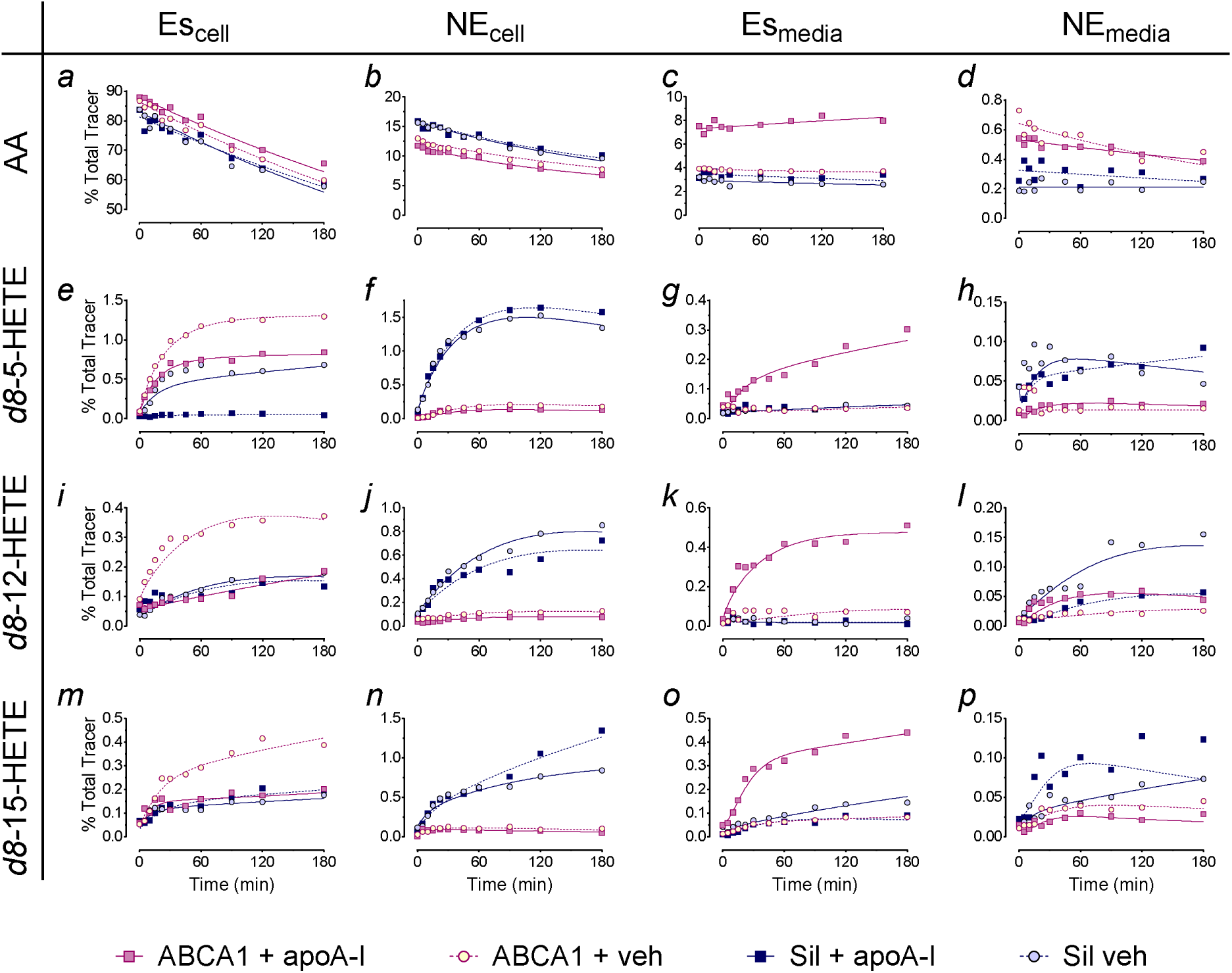
apoA-I/ABCA1-dependent export of HETEs regulates intracellular HETE content. Fit of *d8*-tracer data to the model outlined in Figure 1 in endotoxin-challenged RAW 264.7 macrophages in all 4 treatment conditions. The model evaluates recruitment of AA, intracellular conversion to HETEs, to re-esterification in membrane phospholipids, to export as extracellular HETE. Details of fractional transfer rates (i.e. FTRs) and comparisons between treatments can be found in Table 1. Model details can be found in Supplemental Table 1. The AA and its HETE metabolites are arranged in rows from top to bottom in subpanels: **a-*d*** *= d8*-AA; ***e*-*h*** *= d8*-5-HETE; ***i*-*l*** *= d8*-12-HETE; ***m*-*p*** = *d8*-15-HETE. Each column represents pools from left to right: **Es**_cell_ = cell esterified; **NE**_cell_ = cell non-esterified; **Es**_media_ = media esterified; **NE**_media_ = media non-esterified. *d8*-AA is characterized by: ES_cell_*-d8-AA* disappearance indicating utilization; less rapid NE_cell_-disappearance with lower absolute levels for ABCA1 cells; steady state AA in Es_media_ with much greater concentrations only in the ABCA1 + apoA-I condition; and very low levels of NE_media_ AA which were elevated but declining for cells expressing ABCA1, and low but increasing in Silenced cells. Conversion of AA to 5-HETEs took place in the Es_cell_ pool and conversion to 12-HETE and 15-HETE took place in the NE_cell_ pool. Panels **e, I, m** represent fit for Es_cell_-HETEs, predominantly membrane phospholipids. For *d8*-5-HETE, rapid accumulation takes place for Silenced + veh and ABCA1 + apoA-I condition, however the most rapid accumulation took place in the ABCA1 + veh condition; little accumulation took place in the Silenced + apoA-I condition. in ABCA1 expressing cells, with little dependence on apoA-I. Panels **f, j, n** represent fit for NE_cell_-HETEs and indicate rapid accumulation in the Silenced condition irrespective of aopA-I. Conversely, the presence of ABCA1 was all that was necessary to suppress HETE accumulation. The cell pools together are consistent with an unexpected role of ABCA1 in facilitating re-acylation of HETEs, likely into membrane phospholipids. Additionally, the lack of HETE accumulation in the Es_cell_ pool is also consistent with apoA-I-dependent transfer of accumulating HETEs to the Es_media_. To be consistent with classic HDL lipid efflux, our central hypothesis of HETE-containing phospholipids from cells to HDL would require both ABCA1 and apoA-I. This is evident in Es_media_ (**g**, **k**, **o**) where the *rapid accumulation of esterified d8-HETE occurred only in the ABCA1 + apoA-I condition*. In contrast, cells lacking ABCA1 generally demonstrated more rapid accumulation in the NE_media_ pool (**h**, **l**, **p**) suggesting adaptation to poor Es-efflux capacity by direct diffusion.

**Table 1:** Primary Es- and NE-HETE pool Fractional Transfer Rates and pool transit characteristics.

|  |  |  |  |  | Comparison<sup>A</sup> |  |
| --- | --- | --- | --- | --- | --- | --- |
| Condition | ABCA1 + apoA-I (+/+) | ABCA1 + Veh (+/-) | Sil + apoA-I (-/+) | Sil + Veh<sup>B</sup> (-/-) | +/+ vs +/- | +/+ vs -/+ |
| ***k(12,7); HETE efflux to HDL (Pool Fraction/min)*** |  |  |  |  |  |  |
| 5-HETE | 0.059<br>(0.050, 0.067) | 0.00026<br>(0.00001, 0.00052) | 0.00040<br>(0.00021, 0.0006) | 0.0043<br>(0.0013, 0.0073) | <0.0001 | <0.0001 |
| 12-HETE | 0.234<br>(0.184, 0.284) | 0.005<br>(0.003, 0.007) | 1e-8 | 1e-8 | <0.0001 | <0.0001 |
| 15-HETE | 0.140<br>(0.114, 0.166) | 0.059<br>(0.049, 0.069) | 0.165<br>(0.106, 0.225) | 0.059<br>(0.049, 0.069) | 0.10 | <0.0001 |
| ***NE-HETE Transit Time (min)*** |  |  |  |  |  |  |
| 5-HETE | 15.1 | 15.1 | 0.10 | 0.65 |  |  |
| 12-HETE | 1.4 | 3.8 | 0.04 | 0.04 |  |  |
| 15-HETE | 2.4 | 2.4 | 0.45 | 0.44 |  |  |
| ***NE-HETE Residence Time (min)*** |  |  |  |  |  |  |
| 5-HETE | 18.4 | 19.6 | 11.0 | 7.9 |  |  |
| 12-HETE | 1.5 | 9.2 | 6.9 | 7.1 |  |  |
| 15-HETE | 4.5 | 7.3 | 15.3 | 19.3 |  |  |
| ***NE-HETE Total Transit Number (n)*** |  |  |  |  |  |  |
| 5-HETE | 1.2 | 1.3 | 112 | 12 |  |  |
| 12-HETE | 1.1 | 2.4 | 183 | 189 |  |  |
| 15-HETE | 1.9 | 3.1 | 34 | 44 |  |  |
| ***k(7,10): HETE Re-esterification rate (Pool Fraction/min)*** |  |  |  |  |  |  |
| 5-HETE | 0.066<br>(0.048, 0.084) | 0.066<br>(0.048, 0.084) | 10.1<br>(7.6, 12.6) | 1.42<br>(1.27, 1.57) | e | <0.0001 |
| 12-HETE | 0.656<br>(0.613, 0.700) | 0.228<br>(0.180, 0.276) | 23.6<br>(17.3, 29.9) | 23.6<br>(17.3, 29.9) | <0.0001 | <0.0001 |
| 15-HETE | 0.412<br>(0.229, 0.596) | 0.412<br>(0.229, 0.596) | 2.22<br>(2.21, 2.24) | 2.22<br>(2.21, 2.24) | e | <0.0001 |
| ***k(10,7): HETE PLA<sub>2</sub>-like Removal rate (Pool Fraction/min)*** |  |  |  |  |  |  |
| 5-HETE | 0.013<br>(0.010, 0.016) | 0.013<br>(0.010, 0.016) | 20.8<br>(16.6, 25.1) | 20.8<br>(16.6, 25.1) | e | <0.0001 |
| 12-HETE | 0.025<br>(0.024, 0.027) | 0.025<br>(0.024, 0.027) | 9.53<br>(5.84, 13.22) | 8.72<br>(3.91, 13.54) | e | <0.0001 |
| 15-HETE | 0.143<br>(0.035, 0.252) | 0.143<br>(0.035, 0.252) | 6.80<br>(5.95, 7.66) | 6.80<br>(5.95, 7.66) | e | <0.0001 |

e These parameters are shared
<sup>A</sup> Unadjusted t-test.
<sup>B</sup> The Sil + veh condition was unstable alone and is not used for comparisons. See results.

#### Kinetic characterization of the apoA-I/ABCA1-dependent HETE responses to endotoxin challenge

We evaluated three outcomes for characterizing *d8*-label trafficking: 1) FTR and recycling estimates; 2) area-under-the-curve (AUC) estimates; 3) inter-compartmental *d8*-label flux (*Φ*) estimates. Upon establishing ABCA1/apoA-I dependent transfer of HETEs, we 4) further conducted sensitivity analyses to establish how dependent NE_cell_-HETE concentrations were on the fractional transfer of Es_cell_-HETE to Es_media_ pools; i.e., *k*(12,7).

The FTR estimates relevant to our central hypothesis (i.e., those pertaining to intracellular HETE transport and extracellular efflux) are provided in **Table 1**. FTRs are directly testable features of the model which are independent of tracer or tracee concentrations. Additional features can be calculated from them, including transit times, residence times, and total transits. The efflux of HETEs from the cell Es pool to the proximal Es pool of the nascent media HDL, *(i.e., k(Es-HETE_HDL-p_, Es-HETE_cell_)* described by *k(12,7)*, represents our central mechanistic hypothesis of HDL-mediated impact on intracellular NE HETEs. In the case of 5-HETE and 12-HETE, we found 140-fold and 47-fold greater fractional transfers from Es_cell_ to Es_media_ in the ABCA1/apoA-I condition compared to ABCA1/veh or Silenced/apoA-I (Table 1). Surprisingly, substantial FTRs for 15-HETE were found in all conditions, and apoA-I alone (Sil/apoA-I) was sufficient to impart the same fractional transfer as ABCA1/apoA-I, suggesting that 15-HETE export is partly independent of apoA-I and ABCA1.

A feature of ABCA1/apoA-I dependent export is its potential to reduce the availability of PL-HETEs for recycling. This is evident from transit times and residence times in each compartment. HETE recycling through the Lands cycle^19^ is primarily described by: 1) the removal of NE -HETE by re-esterification into the Es_cell_-HETE pool by ASCL / lysophospholipid acyltranferase (LPLAT)-like activity, [*i.e., k(NE-HETE_cell_, Es-HETE_cell_)*] described by *k(7,10)*; and 2) the transfer of HETEs from the Es_cell_ pool to the NE_cell_ pool by a PLA_2_-like activity *i.e., k(Es-HETE_cell_, NE-HETE _Cell_)* described by *k(10,7)*. Re-esterification was inversely associated with ABCA1 expression: Silenced cells had transfer rates at *k(7,10)* between 5 and 150-fold greater than expressing cells. Similarly, PLA_2_-like release of Es_cell_-HETEs back into the cytosol at *k(10,7)* was rapid in the absence of ABCA1, between 50 and 1,600-fold greater in Silenced cells. The result is much more rapid NE_cell_-HETE transit times (i.e., the average time for a molecule to make a single transit through a compartment) in the absence of ABCA1, however the cumulative residence time is either not shortened (e.g., compare 5-HETE residence times for ABCA1 + apoA-I vs. Sil + apoA-I) or even extended (e.g., compare 15-HETE residence times for ABCA1 + apoA-I vs. Sil + apoA-I). The result is that the primary driver of recycling is the capacity of ABCA1 to modify *k(7,10)* and *k(10,7)* to reduce bi-directional exchange. While the effects of apoA-I are comparatively mild, its absence lengthens each 12-HETE transit and extends the cumulative time a molecule spends in a compartment once entering it (i.e., the residence time) of 12-HETE 6-fold and 15-HETE 1.6-fold. We express recycling as the cumulative transits across a compartment after its initial entrance, hence a value >1 indicates some degree of recycling. The absence of ABCA1 is associated with multiple transits across the NE_cell_-HETE pool, specifically the proximal compartment (pool 10), consistent with increased recycling and a loss of the capacity to exclude Es_cell_-HETEs through PLA_2_-like lipolysis and consequent delivery to the NE_cell_ pool. Recycling of 5-HETE is not appreciably affected by apoA-I, however recycling of both 12-HETE and 15-HETE are increased 2-fold and 1.6-fold respectively in the absence of apoA-I.

Other FTRs of interest are given in **Supplemental Table 3** and include: *k(4,1)* – NE_cell_-AA production from by PLA_2_ on Es_cell_-AA **(ST3A)**; *k(7,1)* – conversion of Es_cell_-AA by *5-LOX* to produce Es_cell_-5-HETE; and *k(10,4)* – conversion of NE_cell_-AA to NE_cell_-12-HETE or NE_cell_-15-HETE by *12/15-LOX* **(ST3B)**. PLA_2_ lipolysis in absence of apoA-I was increased compared to apoA-I + ABCA1, however the absence of ABCA1 resulted in reduced lipolysis. In the case of 5-HETE production, the absence of either ABCA1 or apoA-I resulted in increased conversion rates, however in the case of both 12-HETE and 15-HETE, the absence of ABCA1 had no effect on conversion but the absence of apoA-I resulted in lower conversion.

*<u>AUC estimates</u>* represent the cumulative time-dependent exposure of the pool to labeled metabolites. It is inversely proportional to clearance and is useful because it has lower coefficients of variation compared to FTR estimates. **Supplemental Figure 3** shows results from AUCs, which confirm the findings already reported.

*<u>Estimated flux</u>*; where FTRs represent fractional transfer, Φ describes transfer in absolute terms and is illustrated in **Figure 4**. Net *Φ* is the product of concentration and FTR; [*d8*]*FTR, hence higher concentrations of *d8* could compensate for lower FTRs. *Φ(2,1)_d8_*_-AA_ only occurred in the ABCA1/apoA-I condition **(F4a)**. Substantial Φ*(12,7)*_5-HETE_ occurred only in ABCA1/apoA-I condition **(F4b)**. Similarly, *Φ(12,7)*_12-HETE_ was much greater in the ABCA1/apoA-I condition than any other condition **(F4c).** In contrast, substantial *Φ(12,7)*_15-HETE_ occurred in all treatment conditions **(F4d)**. In the case of the ABCA1/veh condition, while the FTR was low, the effect of ABCA1 to facilitate high Es_cell_*-*15-HETE concentrations overcomes this. In the case of the silenced/apoA-I condition, the transfer rate is comparable to the ABCA1/apoA-I condition is offset by the lack of ABCA1-dependent capacity for accumulating Es_cell_-HETE, thereby limiting net transfer to Es_media_-15-HETE.

**Figure 4:**
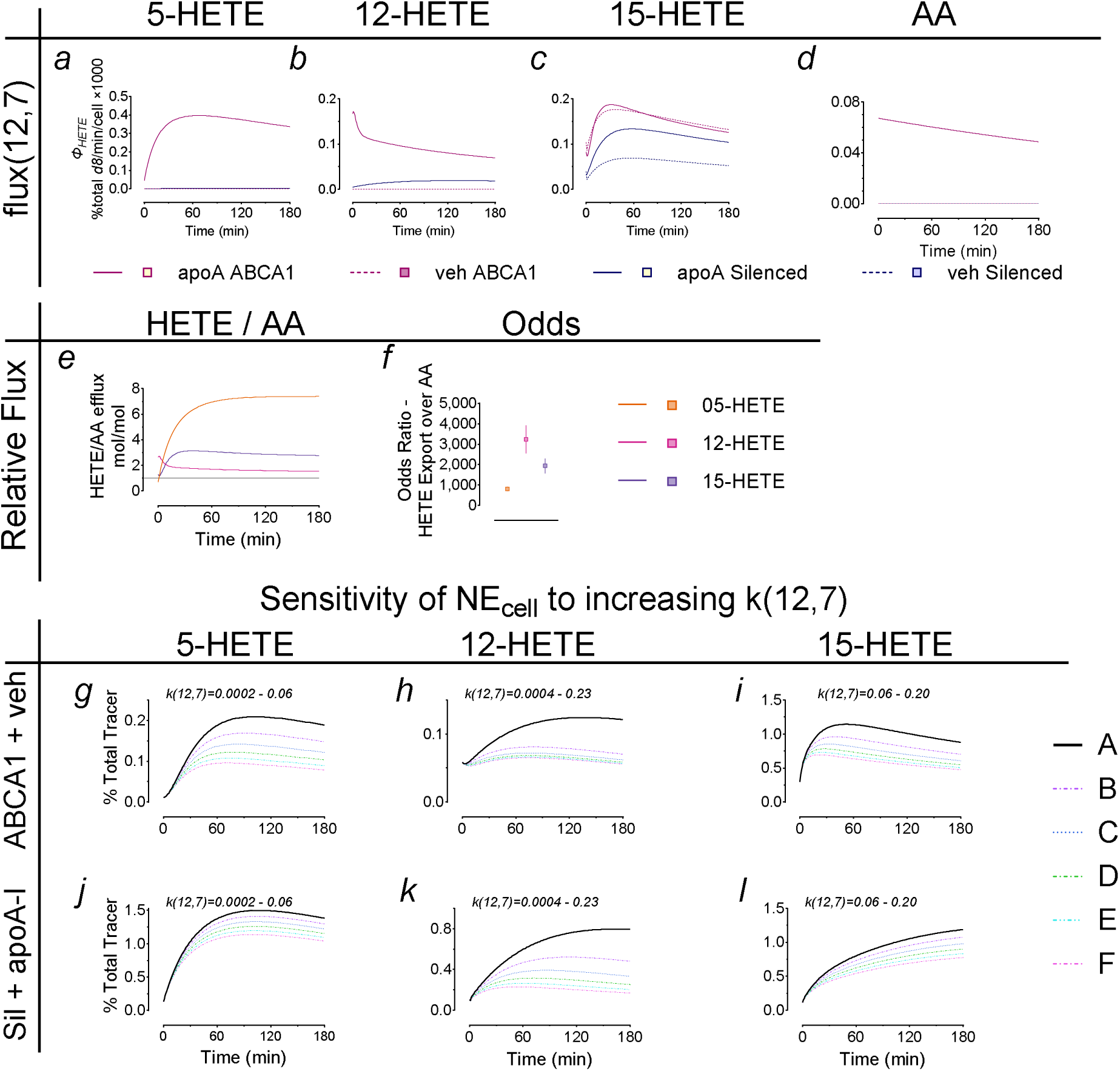
ApoA-I/ABCA1 lipid efflux is selective and regulates NE_cell_ HETEs. *<u>Selective efflux</u>: d8*-HETE efflux was estimated as the percent of total d8-HETE moving from the Es_cell_ pool to the Es_media_ pool per min per cell for each treatment condition and regioisomer **(a-c)** as well as for d8-AA **(d)**, which was only measurable in the apoA-I/ABCA1 condition. For this condition, we were able to estimate the relative efflux of each HETE compared to AA **(e)**. Since Es_cell_ AA is much more abundant i than Es_cell_ HETE, we estimated the proportional odds for HETE efflux relative to AA, which is the ratio of FTR_HETE_/FTR_AA_ **(f)**. *<u>Regulatory effects of efflux</u>*: We conducted sensitivity analyses using the ABCA1 + veh **(g-i)** and Sil + apoA-I **(j-l)** conditions, each which lacked an efflux component and corresponding low HETE flux to Es_media_ (except for 15-HETE). For 5-HETE **(g, j)** and 12-HETE (h, k), we conducted sensitivity analyses as sequentially stepping only k(12,7) from its estimated value to that estimated in the ABCA1 + apoA-I condition in six steps. For 15-HETE (i, l), we simply stepped k(12,7) from 0.06 to 0.2 (since efflux was only partially ABCA1/apoA-I dependent). In all cases, stepping k(12,7) alone was sufficient to reduce the NE_cell_ HETE abundance.

##### HETE efflux is preferential over AA efflux

Another feature of our model is the ability to compare *Φ(2,1)_AA_* to *Φ(12,7)*_HETE_. Es_cell_ *d8*-AA abundance was 900-2200× greater than Es_cell_ *d8*-HETEs. We compared the relative *Φ* of each using their molar ratio of efflux as *Φ(12,7)*_HETE_/*Φ(2,1)*_AA_. Since measurable *Φ(2,1)*_AA_ occurred only in the ABCA1/apoA-I condition, this comparison was only valid in this condition. For all three HETEs the relative *Φ* was >1 indicating that despite its relative scarcity, *Φ*_HETE_ exceeded *Φ*_AA_ **(F4E).** The relative odds for efflux are calculated as *k(12,7)/k(2,1)* **(F4F)** and demonstrate a strong preference for HETE efflux, showing the greatest preference for 12-HETE, and >800× for 5-HETE, the least preferred over AA.

##### Restoration of efflux reduces NE_cell_-HETE abundance

Sensitivity analyses were conducted in both the ABCA1/veh condition, which lacks FTR *k(12,7)* due to lack of apoA-I, and in the Sil/apoA-I condition, which lacks FTR *k(12,7)* due to lack of ABCA1 by constraining all FTRs except *k(12,7)* to their modeled value and increasing *k(12,7)* from its value in the condition to that in ABCA1/apoA-I **(F4g-l)**. In all cases, increasing *k(12,7)*.

#### Correspondence to autooxidation

An alternative source of HETE conversion from AA is autooxidation. To monitor this contribution, we used 9-HETE – because it is not synthesized enzymatically, its only source is non-specific autooxidation. **Supplemental Figure 4** shows the time dependent accumulation of 9-HETE and demonstrates very low rates of appearance in a few select conditions demonstrating that even under strong endotoxin stimulus, HETE production is under enzymatic regulation.

### V: Specific accumulation of 12-HETE in HDL of endotoxin-challenged humans

We have previously reported the response of oxylipins in triglyceride-rich lipoproteins to endotoxin challenge^28^. Here, in an ancillary analysis of HDL in the same 17 subjects challenged with 100 ng/kg LPS, we evaluated the HDL-HETE response under placebo treatment **(Figure 5)**. See **Supplemental Table 3** for baseline subject characteristics. Following endotoxin challenge, HDL-cholesterol declined over 72 hours **(F5A)**, consistent with prior findings^29^. Total HETEs did not decline, but increased marginally in the first hour, followed by a slow decline to baseline over the duration of the observation (168 h). This pattern was reflected in 5-HETE, 15-HETE, and 9-HETE **(F5E)**, the latter of which is a marker for autooxidation since its appearance lacks an enzymatic source. In contrast, we found a sustained time-dependent increase in HDL 12-HETE over the initial 4 hours **(F5C)**, followed by a decline over nearly 72 hours following. Notably, the 15-LOX metabolite of linoleic acid, 13-HODE, declined over the initial four hours following the challenge and remained suppressed for the remaining observational period.

**Figure 5.**
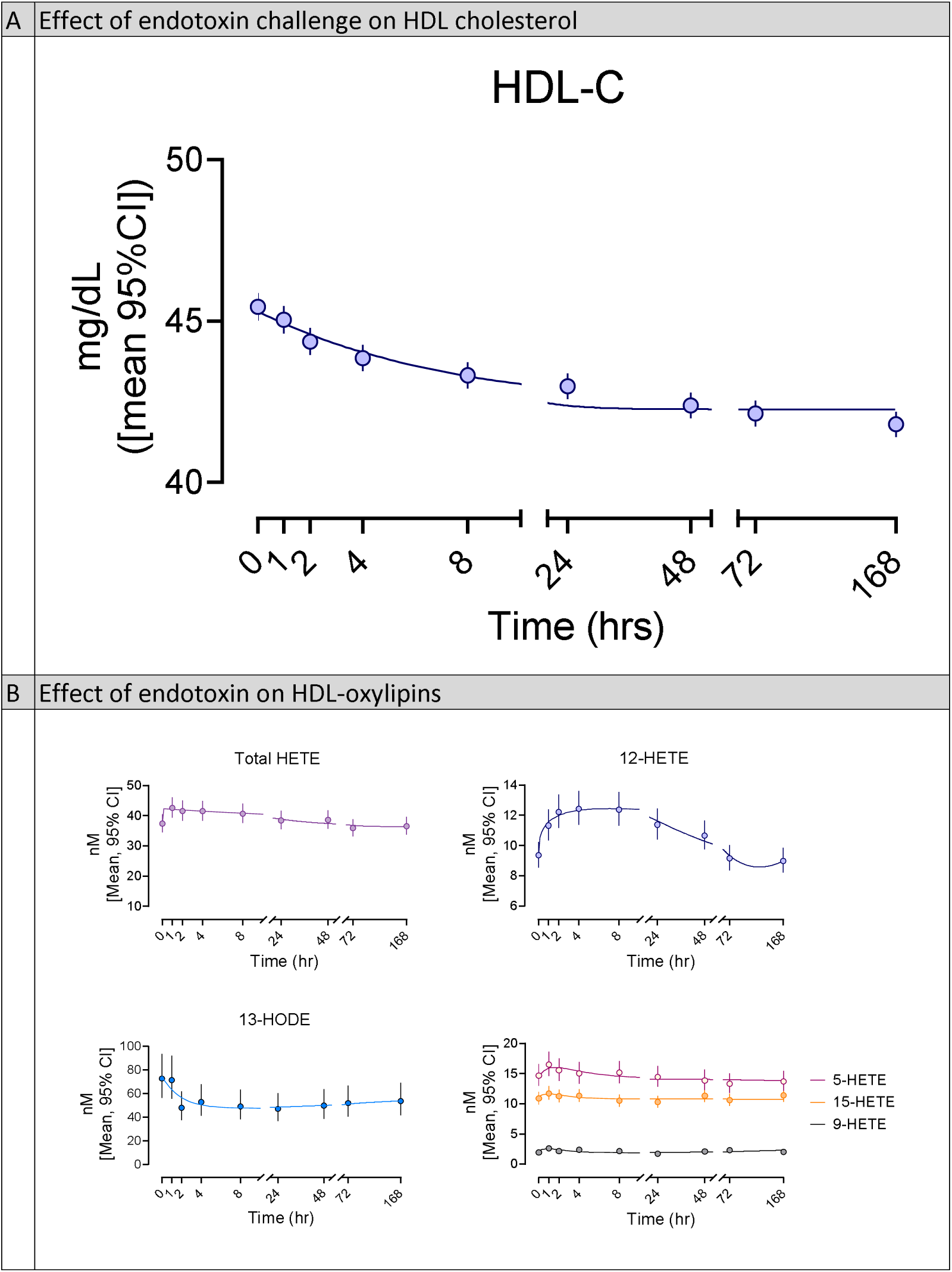
Effects of endotoxin challenge on HDL-C and HETEs. Following endotoxin challenge, there was a time-dependent decline in the mean HDL-C from 45.3 to 42.3 mg/dL, at a rate of 0.136 (0.035, 0.303) pools/hr, which corresponds to a half-life of 5.1 (2.3, 20) hrs **(A)**; total HETEs rose in the first hour and then declined over a 6 nM span for the remaining observational period at 0.036 (0.012, 0.129) pools/hr, which corresponds to a half-life of 19 (5.4, 56) hrs. This pattern was marginally reflected in individual HETEs, such as 5-HETE and 15-HETE and in the autoxidative marker 9-HETE **(B)**. In contrast, 12-HETE followed a substantially different pattern, rising over the first four hours post-challenge, then declining back to nearly baseline over 164 hours. 13-HODE, which is the LA analog of 15-HETE, followed a unique pattern declining initially to the second hour, then undergoing moderate recovery.

## Discussion

While HETEs are important inflammatory mediators, the mechanisms regulating their intracellular fate are not well understood. Our results demonstrate a novel role for ABCA1/apoA-I dependent lipid efflux in regulating intracellular inflammatory signaling. The mechanism has a molecular basis and provides a direct explanation for HDL bioactivity in inflammation. We confirm: 1) Es-HETE efflux is analogous to cholesterol and phospholipid efflux to apoA-I^30,31^ and in the case of nascent HDL is facilitated by ABCA1; 2) HETE efflux acts to regulate the bioactive pool of NE_cell_-HETEs, likely by preventing recycling of HETE; and 3) regulation of NE_cell_-HETE in turn regulates macrophage inflammatory activity, as evidenced by our reporter, MCP-1. We additionally report surprising findings: 4) ABCA1 alone facilitates sequestration of HETEs into the phospholipid pool; 5) a strong preference of ABCA1 for HETEs over AA; and 6) the system has features specific to each HETE which define each HETE’s timeframe and scale of response.

Since 2008 our understanding of HDL metabolism has been challenged, in part due to the failure of multiple clinical trials targeting circulating HDL-cholesterol, including niacin formulations^6^ and CETP inhibitors^7^. The failures resulted in multiple calls for re-evaluation of HDL as a therapeutic^32,33^, with a particular focus on HDL function *en lieu* of static HDL-C levels that continues to this day^34^. Evidence supports the validity of cholesterol efflux capacity as a marker of HDL function in atherosclerotic disease^35^, however a direct molecular link between HDL-mediated lipid exchange and inflammatory function has not emerged. Much work has helped to define the nature of HDL lipid exchange. Segrest et al. demonstrated that ABCA1’s essential activity to facilitate cholesterol export is as a phospholipid translocase^36^. ApoA-I/ABCA1 as an oxylipin efflux system provides a new framework for interpreting extant data. Our work extends a phospholipid-centric understanding of ABCA1 activity by establishing a functional outcome of phospholipid-specific export, independent of cholesterol – modulation of inflammatory activity.

HDL are negatively associated with risk for chronic inflammatory diseases most commonly expressed in risk prediction for chronic inflammatory conditions such as coronary heart disease^37,38^. However, HDL is also important in acute immune function. HDL-C declines in response to severe sepsis^39^ and is inversely associated with inflammatory cytokines TNFα and IL-6^39,40^. Moreover, HDL-C has a U-shaped association with immune response: both low (<31 mg/dL) and high (>100 mg/dL) HDL-C are associated with increased hazards for infectious disease hospitalizations^41^. Here we show that humans responding directly to an endotoxin challenge accumulate 12-HETE in HDL, consistent with a system undergoing regulation of intracellular 12-HETE signaling by HDL-efflux. In our system, we used MCP-1 as a readout for inflammatory activation. MCP-1 is a major chemokine induced by 12-HETE^19^ and is associated with monocyte recruitment to sites of inflammation^42^. Our findings demonstrate that apoA-I and ABCA1 work in concert to reduce MCP-1 expression by limiting the accumulation of NE_cell_-12-HETE, and hence NE-12-HETE is a reliable reporter of macrophage inflammatory response in our system. This novel activity reduces NE_cell_-HETEs, which in turn reduces macrophage inflammatory activity.

Phenomenologically and mechanistically, Es-HETE transfer only occurs when each essential element of lipid transfer from cell to nascent HDL is present, ABCA1 and apoA-I. In conditions lacking the capacity for Es-HETE export (e.g. ABCA1 + veh, Sil + apoA-I), the stepwise restoration of efflux reduced NE_cell_-HETE available for bioactivity. This effect demonstrates the system’s potential for regulatory impact. To the extent elevated HDL-HETEs in humans with metabolic syndrome^43^ are attributable to ABCA1-dependent efflux from cells, this framework implies that these elevations result from enhanced oxylipin efflux activity which would be consistent with restricting or suppressing intracellular inflammatory signaling.

We were surprised by the association of ABCA1 with Es_cell_-HETE in the absence of exogenous apoA-I, which indicates only ABCA1 is required for HETE acylation into membrane phospholipids^44^. Es_cell_-HETE content was much greater in the presence of ABCA1, irrespective of apoA-I. This feature of the system was not straightforwardly obvious, since *k(7,10)* (re-esterification) and *k(10,7)* (PLA_2_-like lipolysis) occur over a longer timeframe in the presence of ABCA1, however the increase in k*(10,7)* due to silencing is roughly 10-fold greater than for *k(7,10)*, Silencing is sufficient to prevent Es_cell_-HETE accumulation. We speculate a leaflet-specific mechanism, since PL-HETEs on the outer-leaflet would not be available for recycling. ABCA1 captures PL on the outer membrane leaflet, however we did not model leaflet specificity.

We were also surprised to find a strong preference for HETE export over AA, which could have important implications for our understanding of HDL function. System specificity implies HETE transport is a primary regulatory feature of the system to regulate intracellular NE-HETEs. Prior to the endotoxin challenge, AA efflux was under steady-state conditions and not responsive to the endotoxin challenge. In contrast, HETE efflux was highly responsive to endotoxin, and highly preferred compared to AA, at unique magnitudes of preference for each HETE, with the greatest preference being for 12-HETE, suggesting partitioning of HETEs regioisomers into subcellular lipid compartments with different ABCA1 access (e.g., nuclear and vesicular membranes vs plasma membranes). Such unique properties provide a basis for understanding observed differences in intracellular HETE signaling, especially in a system where removal, not synthesis rates, drive intracellular abundance.

The demonstration of 12-HETE specific, time-dependent accumulation in humans responding to the same stimulus as the model macrophages supports our in vitro findings, however the lack of 5-HETE accumulation in humans suggest differences between the in vitro and in vivo environment (such as PON1-mediated hydrolase activity on delta-lactones of 5-HETE^45^), species specific difference or differences attributable to the stronger endotoxin stimulus in vitro. We have previously shown the lack of Ffar4, a receptor responsible for activation of cPLA_2α_, results in substantial changes in circulating HDL oxylipin content whether derived from mice with experimental heart failure or metabolic disease, in particular the accumulation of 12-HETE relative to 18-HEPE, an EPA-derived alcohol^46,47^. The lack of 12-HETE in the HDL of animals with disrupted PLA_2_-signaling demonstrates the potential for specific signaling deficits to be evident in circulating HDL oxylipin content.

Ultimately, we confirm and/or demonstrate six important features of the described apoA-I/ABCA1-dependent intracellular HETE regulatory system: 1) intracellular NE-HETEs regulate inflammatory outcomes; 2) apoA-I and ABCA1 are essential elements of the transfer of Es-HETE from cells to HDL; 3) the activities of apoA-I and ABCA1 contribute to the regulation of intracellular NE-HETEs; 4) HETE regioisomers have unique efflux properties impacting differences in their duration, transit, and recycling in the intracellular NE-HETE pool; 5) the efflux system appears highly preferential to HETEs over AA, suggesting HETE efflux is a principal function of apoA-I/ABCA1-dependent efflux explored here.

### Strengths and limitations

This experiment leverages well characterized systems in HDL biology, inflammation, and intracellular cytokine production to validate a novel regulatory system. The tracer analytics provides for strong inferences to the HETE origins and trafficking, and the compartmental models allow for mechanistic insight into the system features. Finally, observation of the expected response to the identical challenge in humans provides *in vivo* support for more detailed findings in vitro.

This study is limited by its exploratory and novel nature. While we establish how the system works and provide explanatory power for the overlap of inflammation and HDL biology, the findings require broader validation before applying them to specific systems. Further, we interpret our finding within the framework of the canonical pathway for PLA_2_-mediated signaling, in which the cytosolic NE-HETEs are bioactive. Emerging evidence suggests 12-HETE has bioactivity as a component of phospholipids^48^. Since we found that apoA-I and ABCA1 also have profound consequences for cell Es-HETEs, re-interpretation is warranted as new evidence emerges.

In conclusion, we report a molecular explanation for HDL regulation of inflammation: reduction of pro-inflammatory NE_cell_-HETE signaling by removal of Es_cell_-HETE from cell membranes to nascent HDL. This finding provides an explanation for the associations of HDL with inflammatory metabolic disorders. Therefore, the apoA-I/ABCA1 system which mediates cholesterol and phospholipid efflux also participates in the efflux of esterified HETEs contributing to the regulation of the bioactive NE-HETEs in intracellular oxylipin signaling. This novel system in which HDL-lipid efflux participates in regulation of intracellular signaling, provides powerful molecular explanations for the association of HDL with anti-inflammatory activity. This insight should be useful for better diagnostic and interventional efforts.

We expect activities of PLA_2_ and LOX participate in the acute regulation of intracellular NE_cell_-HETEs, however in our test conditions they did not contribute strongly to establishing the rate of time-dependent NE_cell_-HETE accumulation. Instead, we found intracellular trafficking post-HETE conversion, which is specific to each HETE, could explain the observed differences in NE_cell_-HETE concentrations.

In addition, we identified unique features of 15-HETE efflux that were not entirely dependent on either apoA-I or ABCA1 suggesting other mediators of HDL-lipid transfer such as ABCG1 or SR-B1 expression ^32^ may also participate in the regulation of 15-HETE efflux.

## Research Design and Methods

### Cell Culture and siRNA Transfection

Two separate cell culture experiments were conducted, the first to verify the inflammatory model in using 4 different timepoints (0 min, 60 min, 120 min, and 180 min) and the second utilizing tracers AA-d8 with 11 different timepoints (0 min, 5 min, 10 min, 15 min, 22.5 min, 30 min, 45 min, 60 min, 90 min, 120 min, and 180 min). RAW 264.7 cells were grown in Dulbecco’s modified eagle medium (DMEM) containing 10% (v/v) fetal bovine serum (FBS) (heat-inactivated) with 1% (v/v) penicillin/streptomycin at 37°C in 5% CO2. Cells were plated and grown to recommended density per Lipofectamine™ 3000 (Invitrogen) protocol. Briefly, 5 µg of ABCA1 siRNAs (511GUGGCCUGGCCUCUCUUUAddT311), positive control GAPDH siRNA, and scrambled negative control siRNAs were diluted in separate vials of Opti-MEM and Lipofectamine™ and incubated at room temperature. Mixture was then added to cell culture wells and incubated for 2 days. Cells were then supplemented with AA-d8 for 8 hours prior to treatment, washed twice, and replaced with FBS free media containing Opti-MEM Reduced Serum with 1% (v/v) penicillin/streptomycin treated with 8-Br-cAMP (50uM), the LXR agonist TO901317 (10uM), and +/- ApoA-1 (40ug/mL) for 2 hours. Cells were then treated with 100 ng/mL of LPS after which 1 mL of media was collected and snap frozen. Cells were washed twice with PBS, scraped, with a small portion stained to evaluate viability with the remainder snap frozen and stored at -80°C until extraction and analysis.

### Cell Culture and siRNA Transfection

We tested the hypothesis that apoA-I/ABCA1-mediated efflux of Es_cell_-HETEs to nascent HDL limits cellular HETE accumulation by reducing the available HETE pool for recycling back to the NE pool. A compartmental model was created using the time-dependent changes in d8-AA and -HETEs through the NE and Es pools in cells and media based on the conceptual and experimental frameworks described in **Figure 2a** and **2b**, respectively.

### RNA Extraction, Reverse-Transcription, and qRT-PCR

Total RNA was extracted from one six-well plate using TRIzol Reagent (Invitrogen) and for each treatment group and time. One microgram of RNA was reverse transcribed into cDNA using Superscript II reverse transcriptase (Invitrogen). The mRNA concentrations were determined by qRT-PCR using SYBR Select Master Mix with primer sequences specific to mouse ABCA1. Glyceraldehyde-3-phosphate dehydrogenase (GAPDH) was used as a reference and qPCR conditions were set to 40 cycles of 95°C for 20 s, 95°C for 30 s, and 60°C for 30. Measurement of mRNA expression was calculated by the ΔCt method using GAPDH as an internal control. Primers used for qRT-PCR are shown in **Supplemental Table 4**.

### Western Blot Analysis

Cells were lysed in 1% NP-40 buffer with added protease inhibitors. The protein concentration was measured using a Bradford Protein Assay kit (Bio-RAD). 40 µg of lysate was added to 4-15% Tris-HCL SDS gel (Bio-RAD) in sample buffer and then transferred to nitrocellulose membrane at 30V overnight. Membranes were blocked in 5% milk in TBS-T for 2 hours and then incubated with primary antibodies against GAPDH (NB100-56875, 1:1000) and ABCA1 (NB400-105, 1:1000) for 1 hour, washed, and blotted with horseradish peroxidase (HRP)-conjugated IgG. Immunoreactive bands were detected using an ECL detection system (Amersham Bioscience).

### MSD and MCP-1 ELISA

Ten plasma cytokines (IL-9, MCP-1, IL-33, IL-27, IL-15, MIP-1α, IL-6, MIP-2, IL-1β, IL-10) were measured using a V-PLEX Custom Kit from Meso Scale Discovery (MSD, Gaithersburg, MD, USA). All assays were performed in duplicate. Analyses were done using a QuickPlex SQ 120 instrument (MSD) and DISCOVERY WORKBENCH® 4.0 software. MCP-1 ELISA kits and required reagents were used to detect and measure MCP-1 concentrations in the single cell group experiments in accordance with manufacturer’s instruction (Thermo Fisher Scientific).

### Subject Selection and Study Design

This is an ancillary study from a crossover design trial using P-OM3 treatment to measure inflammatory response difference after endotoxin challenge^28^. Subjects were 20–45-year-old healthy males (N=20). Inclusion criteria required a resting heart rate > 55 bpm, body mass index of 20-30 kg/m^2^, and a fasting LDL-C < 160 mg/dL and 1 serving or less of oily fish per week. Subjects were randomized to placebo and P-OM3 treatment for 8-12 weeks. Following the treatment period was a wash-out of 8 weeks followed with the opposite treatment for 8-12 weeks. 4 capsules of P-OM3 were taken daily totaling 3.4 g/d of EPA + DHA (Pronova Biopharma; Oslo, Norway) or the placebo consisting of 4 capsules taken daily with an olive oil composition. Three subjects from the parent trial denied permission for their samples to be used in future research bringing the sample size for this study to 17. All study protocols were reviewed and approved by the Penn State Institutional Review Board and informed consent was obtained (NCT01813110).

### Endotoxin Challenge

After each 8–12-week treatments, subjects were instructed to fast for 12 hours and avoid alcohol, strenuous exercise, and anti-inflammatory medications for 48 hours prior arrive at the clinical research center (CRC) at Penn State University Park campus. Subjects were administered a low-dose endotoxin challenge calculated as 0.6 ng of LPS/kg body weight and followed with blood sample collections at baseline, 1, 2, 4, and 8 hours. Participants remained at the CRC for an additional 8 hours for monitoring and to receive continuous IV saline. After evaluation by a nurse for stable vital signs, IV catheters were removed, and subjects were allowed to leave the CRC. Subjects were instructed to avoid alcohol, anti-inflammatory medication, limit strenuous activity, and notify the clinic if any symptoms occurred. Subjects returned to the CRC for additional blood sample collection 24, 48, 72, and 168 hours post endotoxin challenge but were not required to be fasted. EDTA tubes were used to collect and centrifuge blood samples to remove red blood cells from plasma, and all samples were stored at -80°C until analysis.

### Clinical Measurements

Plasma total cholesterol, HDL-C, LDL-C, triglycerides (TG), glucose, and insulin were measured at initial screening and at both baseline time points prior to endotoxin challenge being given by Quest Diagnostics.

### Lipoprotein Separation

EDTA plasma samples were used for lipoprotein fraction separation using density ultracentrifugation, methods previously described by Edelstein and Scanu^49^. Plasma was diluted with 1.0063 g/mL sodium chloride saline solution with BHT and EDTA. Plasma sample volumes were all brought to the same level using solution to fill 4.7 mL Optiseal centrifuge tubes (Beckman Coulter; Brea, CA). Samples were centrifuged at 45,000 RPM for 19 hours at 4°C in a XL-70 ultracentrifuge (Beckman Coulter). Centrifuged sample tubes were then sliced at the VLDL band removing the top layer and transferring the remainder to a new Optiseal centrifuge tube. Addition of 1.67 mL 1.1816 g/mL solution of NaCl and NaBr was added to the tubes, capped, and centrifuged at 45,000 RPM for 19 hours at 4°C. Tubes were again sliced at the LDL band removing the top layer and transferring the remaining volume to a new Optiseal centrifuge tube. Addition of 1.67 mL 1.4744 g/mL solution of NaBr was added to the tubes, capped, and centrifuged at 45,000 RPM for 19 hours at 4°C. Tubes were sliced at the HDL band removing the top layer to be stored at -80°C for further analysis.

### HDL-C Colorimetric Measurement

HDL samples were thawed on ice and reagents prepared according to manufacture protocol. In 2 mL test tubes, 0.2 mL of HDL samples and 0.2 mL of the precipitating reagent were added, vortexed, and allowed to stand at room temperature for 10 minutes. The samples were then centrifuged for 10 minutes at 3000rpm and the top 12.5 µL taken and added to the 96 well plate with an addition of 750 mL of the color solution. Standards were made according to the calibration procedure given by the manufacture (Wako, Lexington, MA). The plates were incubated at 37°°C for 5 minutes and HDL-C was measured at 600nm wavelengths.

### Oxylipin Extraction

HDL samples were thawed on ice and extracted using a modified Smedes protocol^50^. 100 µL of media, solution containing cells, and solution containing HDL was spiked with BHT/EDTA (0.2 mg/mL), one deuterated oxylipin surrogates (20 µL of 1000 nM concentration) and subjected to liquid-liquid extraction to isolate lipid content. Samples were split in two, a non-hydrolyzed portion for free oxylipin measurement and a hydrolyzed portion using 0.1 M methanolic sodium hydroxide for total oxylipin measurement and further purified by solid phase extraction using Chromabond HLB sorbent columns. Oxylipins were eluted with 0.5 mL of methanol with 0.1% acetic acid and 1 mL of ethyl acetate, dried under nitrogen stream, and reconstituted with 100 nM 1-cyclohexyluriedo-3-dodecanoic acid (CUDA) in methanol acetonitrile (1:1).

### LCMS Analysis

Samples were analyzed by liquid chromatography using a Waters Acquity UPLC coupled to Waters Xevo triple quadrupole mass spectrometer equipped with electrospray ionization source (Waters, Milford, MA). A volume of 5 µL was injected and separation was performed using a CORTECS UPLC C18 2.1 x 100 mm with 1.6 µM particle size column. Flow rate was set at 0.5 mL/min and consisted of a gradient run using water with 0.1% acetic acid (Solvent A) and acetonitrile isopropanol, 90:10 (Solvent B) for 15 minutes (0-12 min from 25% B to 95% B, 12-12.5 min 95% B, 12.5-15 min 25% B). Electrospray ionization operated in negative ion mode with capillary set at 2.7 kV, desolvation temperature set at 600 °C, and source temp set to 150°C. Optimization of oxylipin MRM transitions were identified by direct injection of pure standards onto the mass spectrometer and using cone voltage and collision energy ramps to optimize detection and most prevalent daughter fragments **(Supplementary Table 4)**.

Calibration curves were generated prior to each run using standards for each oxylipin. Peak detection and integrations were achieved through Target Lynx (Waters, Milford, MA) and each peak inspected for accuracy and corrected when needed. Representative chromatogram traces are shown in **Supplemental Figure 6**.

### Statistical Methods

Kinetic models were built using SAAM II v2.3.3, Nanomath LLC. Structural equation models were developed using AMOS v29.0. Static curve fitting was accomplished using GraphPad Prism 10.1.2, and mixed model ANOVAs were used to evaluate the time-dependent response to endotoxin challenge using JMP 17.0.0. Each testing environment emphasizes unique outcomes and goals.

For compartmental modeling using SAAM II, model development followed the methods and recommendations of Wastney et al^51^ and the simultaneous fit of all model data, which maximizes the value of information on the system. Evaluative criteria were 1) consistency with the biological properties of the system, and 2) mathematically acceptable fit. Unique features of our testing environment were the opportunity to share FTRs between conditions and between HETE systems. We worked from an initial construct with single compartments corresponding to each tracer pool measured, four for *d8*-AA and four for each *d8*-HETEs, with FTRs representing the Lands’ cycle for *d8*-AA, and identical recycling of HETEs. These FTRs were supplemented with transfers of cell-Es HETEs and NE HETEs to their respective media pool. Mathematical considerations include a consistent and unique solution with minimal error; an acceptable mathematical fit was considered to minimize the objective function, have a fractional standard deviation <0.5, no parameter correlations <-0.8 or >0.8, a random residual distribution, and a small sum of squared residuals between compartments. Models were modified iteratively by examination of the fit to data in each pool, examination of the correlation matrix, addition of shared parameters to facilitate unique solutions, and addition of compartments and associated FTRs. Parsimony was improved by forcing FTRs to be equal across conditions with care taken to consider the factorial nature of the experiment and the comparative aims of the experiment.

Unnecessary FTRs were fixed to 1e-8 to preserve matrix invertibility. Satisfactory fit required the addition of four additional compartments: one to describe the behavior of intracellular NE-HETE (HETE_Othercell_11), two to describe cell Es-HETE (Es-HETE_Cell-2_8; Es-HETE_Cell-3_9), and one to describe extracellular Es-HETE (Es-HETE_HDL-d_13). Their addition improved conformity to observed data and satisfy the biological likelihood that: 1) cell NE-HETE are sub-compartmented within specific organelles, 2) cell Es-HETEs are likely to be re-esterified into multiple organelles such as the mitochondria^52^ in addition to specific leaflets of the plasma membrane, and 3) extracellular Es-HETEs are likely to be sorted in HDL in the same manner as phospholipids are, e.g. LCAT mediated transfer from phospholipids to cholesterol esters. Unique solutions were ensured by evaluation of the correlation matrix and sharing parameters. FTRs are directly estimated from the compartmental model and represented conventionally as k(*i*,*j*) where *i* is the compartment transferred to, and *j* is the compartment of origin. Flux at time t (*Φ*_t_) is calculated as a product of the FTR and the *d8*-tracer concentration at time t and is represented conventionally as *Φ*(*i*,*j*). AUC is calculated using the Rosenbrock integration method for area under the curve.

For mediation analysis by pathway modeling using AMOS 29.0.0, structural equation models were developed following the methods and recommendations of Hayes^53^ using Time, Time^2^, Time^3^, ABCA1, apoA-I, and ABCA1×apoA-I as predictors of MCP-1, and NE-12-HETE as the mediator. A cubic polynomial function was sufficient to account for the non-linear relationship of time to 12-HETE and MCP-1, which in turn allowed for estimation of the direct effect of all predictors on MCP-1, and the mediated effects of all predictors on MCP-1 by 12-HETE. Parameter elimination was facilitated by the *specification search* feature and model fit was confirmed by the Χ^2^ test, the goodness of fit index (GFI), and other supporting indices. Post-hoc differences were evaluated with user defined equations.

Graph Pad Prism was used to describe the time-dependent changes in total HETEs and in the mean human total HDL HETEs. The aim of the illustrations is descriptive, hence the goal of fit was to find the best model having optimally shared parameters across all datasets using BIC as a selection criteria for best description. No statistical comparisons are made, however for the human data results the associated slopes are reported to allow a succinct description of the trends.

Time dependent changes in human HDL lipids were tested for with JMP 17.0.0 using mixed models adjusted for lipid concentration prior to endotoxin challenge. The specific time-dependent repeated covariance structure was selected to minimize BIC.

## Supporting information

Supplemental Results and Figures

## Acknowledgements

We acknowledge the following collaborators who conducted the parent study of human males challenged with LPS^28^: Penny Kris-Etherton, Chesney Richter, Ann Skulas-Ray, Michael Flock, and Gordon Jensen.

