## Supplemental Results and Figures for "ABCA1 and apoA-I dependent 12-hydroxyeicosatetraenoic acid efflux regulates macrophage inflammatory signaling"

**Supplemental Data**

**Supplemental Figure 1 – Validation of treatment conditions**

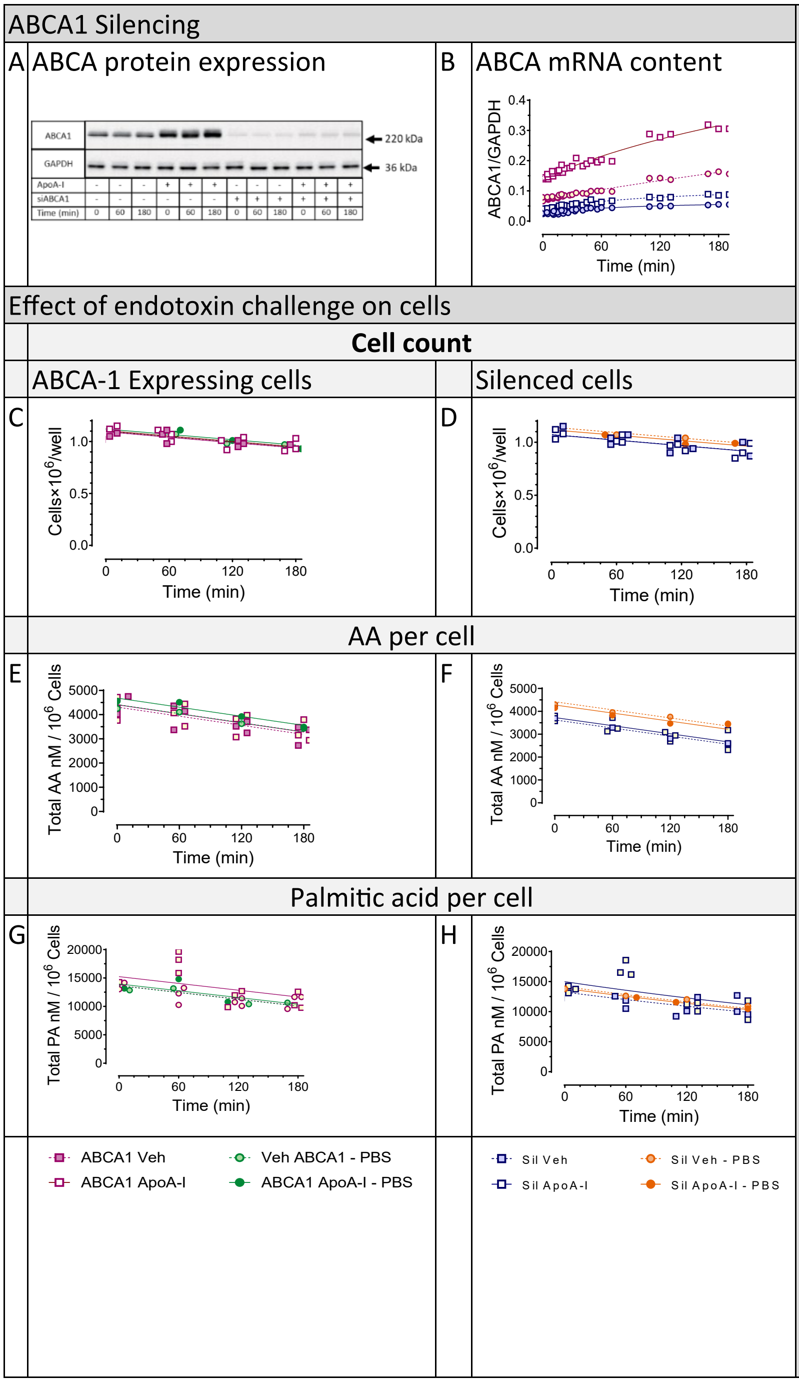

Cells were lysed and purified prior to blot with ABCA1 antibody. GAPDH was used as a control. Bands were noted at 220 kDa for ABCA1 in the ABCA1 expressing cells with increased banding in the ABCA1 treated with ApoA-I. Silenced ABCA1 cells showed significant silencing of ABCA1 protein with slight rescuing in ApoA-I added cells. The four treatment groups: Veh/ABCA1, ApoA-I/ABCA1, Veh/Sil, and ApoA-I/Sil. **A)** Silence mRNA for ABCA1 sequence used **B)** Western blot of ABCA1 for time points 0 minutes, 60 minutes, and 180 minutes for all 4 treatment groups **C)** ABCA1 mRNA expression was measured by qRT-PCR and normalized to GAPDH, in each treatment group after 2 hours of ApoA-I supplementation and prior to LPS. All treatment groups (ApoA-I/ABCA1, Veh/Sil, and ApoA-I/Sil) showed significant differences in ABCA1 expression compared to control (Veh/ABCA1) **D)** Cell counts for each treatment group over time. **E)** GAPDH totals per 1x10^6 cells **F)** Total palmitic acid (PA) concentrations for each treatment group over time per cells **G)** Total PA per GAPDH **H)** Total AA per cells **I)** Total AA per GAPDH **J)** PCR measured ABCA1, normalized to GAPDH, in each treatment group after 2 hours ApoA-I supplementation and no LPS. All treatment groups (ApoA-I/ABCA1, Veh/Sil, and ApoA-I/Sil) showed significant differences in ABCA1 expression compared to control (Veh/ABCA1) **K)** Total ABCA1 per cells for all 4 treatment groups over time. Data were compared by Tukey HSD post hoc analysis, $p<0.001$

**Supplemental Table 1A: Model summary**

| | |
|---|---|
| Independent samples[A] | 132 |
| Pool measurements[A] | 528 |
| *d8*-analytes measured[A] | 2112 |
| N[B] | 696 |
| Estimated Parameters | 90 |
| Set equal[C] | 58 |
| Linearly derived[D] | 16 |
| Total Derived[E] | 74 |
| Total transfers | 164 |
| dF | 532 |
| Total objective[F] | -15.1902 |
| BIC[G] | -5.96838 |

[A] a total of 132 independent cultures of RAW 264.7 cells were included, representing triplicate measurements at each timepoint × condition included in the models. In sample analysis, four pools were extracted from each: $Es_{cell}$, $NE_{cell}$, $Es_{media}$, and $NE_{media}$. Hence, 528 pool measurements were made. Each pool analysis yielded quantitation on four traced analytes (*d8*-AA, *d8*-5-HETE, *d8*-12HETE, and *d8*-15-HETE) and four parent analytes (AA, 5-HETE, 12HETE, and 15-HETE).

[B] Total number of data points in the model following triplicate averaging and accounting for failed collection of *d8*-AA in $Es_{media}$ and $NE_{media}$ in all conditions at 45 minutes.

[C] Total number of shared parameters across conditions or HETEs.

[D] Linearly derived parameters were estimated when correlation (ρ) with other parameters exceeded 0.9. The linear relationship was estimated from three successive solutions where one parameter was solved while the other was fixed.

[E] This is the total number of FTRs derived from shared or linearly estimated parameters. They are not additive since some parameters were shared multiple times.

[F] This is the objective function SAAM-II minimizes to achieve a best fit. Individual objectives are available below in Supplemental Table 1B.

[G] Bayesian information criteria provided an additional tool for comparing models.

**Supplemental Table 1B: Model summary of objective and scaled data variance by sampled pool.**

| Condition | Pool | Objective | | | | Scaled Data Variance | | | |
|---|---|---|---|---|---|---|---|---|---|
| | | AA | 05HT | 12HT | 15HT | AA | 05HT | 12HT | 15HT |
| ABCA1 + apoA-I | $Es_{cell}$ | -0.11 | -0.24 | -0.27 | -0.24 | 0.06 | 0.33 | 1.54 | 3.84 |
| ABCA1 + veh | $Es_{cell}$ | -0.12 | -0.24 | -0.24 | -0.24 | 0.03 | 0.17 | 1.38 | 2.30 |
| Sil + apoA-I | $Es_{cell}$ | -0.10 | -0.20 | -0.27 | -0.26 | 0.09 | 7.51 | 1.27 | 1.35 |
| Sil + veh | $Es_{cell}$ | -0.11 | -0.27 | -0.24 | -0.26 | 0.07 | 6.41 | 8.30 | 1.06 |
| ABCA1 + apoA-I | $NE_{cell}$ | -0.17 | -0.25 | -0.31 | -0.28 | 0.08 | 5.45 | 0.31 | 1.45 |
| ABCA1 + veh | $NE_{cell}$ | -0.16 | -0.27 | -0.29 | -0.26 | 0.14 | 1.45 | 0.45 | 2.09 |
| Sil + apoA-I | $NE_{cell}$ | -0.16 | -0.20 | -0.24 | -0.20 | 0.08 | 1.13 | 0.49 | 5.19 |
| Sil + veh | $NE_{cell}$ | -0.14 | -0.22 | -0.22 | -0.22 | 0.20 | 0.38 | 2.60 | 1.58 |
| ABCA1 + apoA-I | $Es_{media}$ | -0.15 | -0.24 | -0.22 | -0.25 | 0.19 | 3.88 | 3.58 | 1.25 |
| ABCA1 + veh | $Es_{media}$ | -0.18 | -0.28 | -0.24 | -0.32 | 0.06 | 4.68 | 21.43 | 0.19 |
| Sil + apoA-I | $Es_{media}$ | -0.16 | -0.30 | -0.27 | -0.27 | 0.60 | 2.26 | 14.22 | 2.61 |
| Sil + veh | $Es_{media}$ | -0.15 | -0.28 | -0.27 | -0.26 | 0.57 | 6.93 | 31.22 | 12.90 |
| ABCA1 + apoA-I | $NE_{media}$ | -0.22 | -0.31 | -0.27 | -0.30 | 0.20 | 2.93 | 12.86 | 3.88 |
| ABCA1 + veh | $NE_{media}$ | -0.19 | -0.28 | -0.30 | -0.29 | 1.46 | 17.78 | 4.79 | 2.94 |
| Sil + apoA-I | $NE_{media}$ | -0.21 | -0.25 | -0.26 | -0.28 | 2.40 | 10.26 | 4.47 | 4.20 |
| Sil + veh | $NE_{media}$ | -0.19 | -0.28 | -0.30 | -0.25 | 4.73 | 1.57 | 2.18 | 9.45 |

The objective (left) and scaled data variance (right) are represented as heatmaps to evaluate factors driving model fit as well as overall fit of each pool.

**Supplemental Figure 2.**

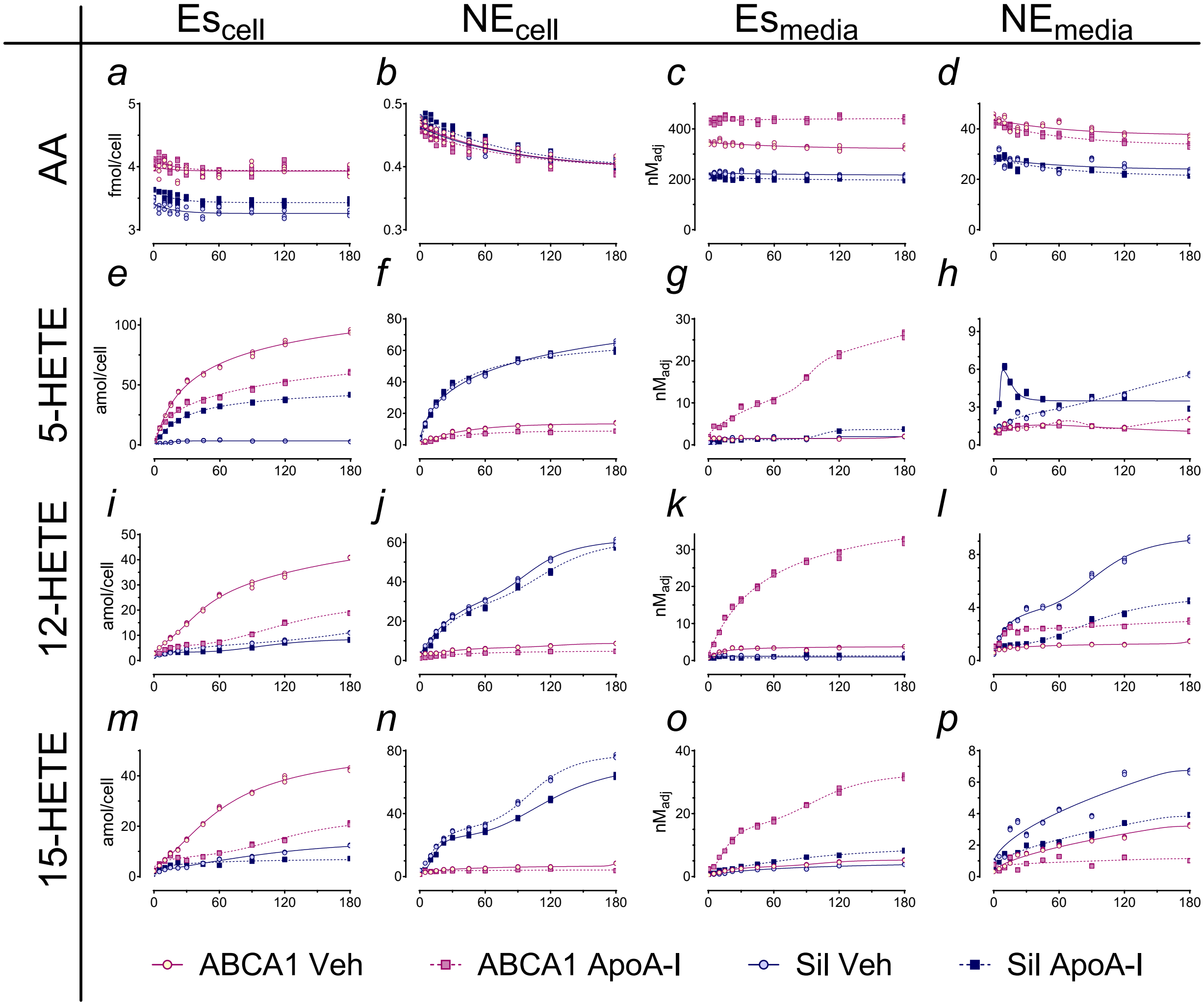

**Supplemental Figure 2: HETE accumulation in each pool.** Total HETE concentration in each pool in endotoxin-challenged RAW 264.7 macrophages in all 4 treatment conditions. The cumulative concentration (labeled plus unlabeled) in each pool conforming to **Fig 2** from recruitment of AA, intracellular conversion to HETEs, to re-esterification in membrane phospholipids, to export as extracellular HETE. The AA and its HETE metabolites are arranged in rows from top to bottom in subpanels: **a-*d*** = AA; ***e-h*** = 5-HETE; ***i-l*** = 12-HETE; ***m-p*** = 15-HETE. Each column represents pools from left to right: **$Es_{cell}$** = cell esterified; **$NE_{cell}$** = cell non-esterified; **$Es_{media}$** = media esterified; **$NE_{media}$** = media non-esterified. Graph Pad Prism was used to find nominally best fit static models, selecting from their dose-response suite of equations and using parameter sharing and AICc to identify a nominal description of the accumulation, consistent with a system of complex interchange between pools and compartments. Panels **e, i, m** represent fit for $Es_{cell}$-HETEs, predominantly in membrane phospholipids. Panels **f, j ,n** represent fit for $NE_{cell}$-HETEs. Panels **g**, **k**, **o** represent accumulation in the $Es_{media}$ pool, i.e. HETEs in HDL phospholipids, and finally accumulation in the $NE_{media}$ pool (**h**, **l**, **p**). The HETE concentrations and accumulation rates are well explained by the compartmental model in **Figure 3** and **Table 1**.

**Supplemental Table 2: Additional Fractional Transfer Rates**

**Supplemental Table 2A**

| Condition | ABCA1 + apoA-I (+/+) | ABCA1 + Veh (+/-) | Sil + apoA-I (-/+) | Sil + Veh[B] (-/-) | Comparison[A] +/+ vs +/- | Comparison[A] +/+ vs -/+ |
|---|---|---|---|---|---|---|
| ***k(4,1); $PLA_2$ activity on AA (Pool Fraction/min)*** | | | | | | |
| AA | 0.00127 (0.00114, 0.00139) | 0.00146 (0.00134, 0.00158) | 0.00098 (0.00075, 0.0012) | 0.00128 (0.00103, 0.00152) | <0.0001 | <0.0001 |

**Supplemental Table 2B**

| Condition | ABCA1 + apoA-I (+/+) | ABCA1 + Veh (+/-) | Sil + apoA-I (-/+) | Sil + Veh[B] (-/-) | Comparison[A] +/+ vs +/- | Comparison[A] +/+ vs -/+ |
|---|---|---|---|---|---|---|
| ***AA conversion to HETE (Pool Fraction/min)*** **$Es_{cell}$ AA to $Es_{cell}$ 5-HETE – *k(7,1)*** | | | | | | |
| 5-HETE | 0.000536 (0.000483, 0.000589) | 0.000658 (0.000614, 0.000703) | 0.000960 (0.00079, 0.00113) | 0.000827 (0.000726, 0.000928) | <0.0001 | <0.0001 |
| ***AA conversion to HETE (Pool Fraction/min)*** **$NE_{cell}$ AA to $NE_{cell}$ 5-HETE – *k(10,4)*** | | | | | | |
| 12-HETE | 0.00121 (0.00103, 0.00139) | 0.000785 (0.000749, 0.000821) | 0.00110 (0.00103, 0.00116) | 0.000822 (0.000658, 0.000986) | <0.0001 | 0.12 |
| 15-HETE | 0.00198 (0.00174, 0.00223) | 0.001510 (0.00137, 0.00164) | 0.00170 (0.00134, 0.00206) | 0.00167 (0.00137, 0.00196) | <0.0001 | 0.10 |

FTR of label from the cell esterified pool to the NE pool **(3A)** representing $PLA_2$ activity on AA [*k(4,1)*], was lowest in apoA-I/ABCA1 cells; the lack of either apoA-I or ABCA1 was associated with greater FTRs. FTRs from NE-AA to NE-HETE, *k(10,4)* represents LOX activity **(3B)**. In mouse-derived RAW-264.7 cells, conversion of AA to 5-HETE is mediated by 5-LOX, however both 12-HETE and 15-HETE are produced from a single 12/15-LOX. Conversion of AA to both 12-HETE and 15-HETE was more rapid however in the absence of apoA-I, however in the absence of ABCA1 there was no difference.

[A] unadjusted t-tests

**Supplemental Figure 3: Total area-under the curve exposure of pools to *d8*-label.**

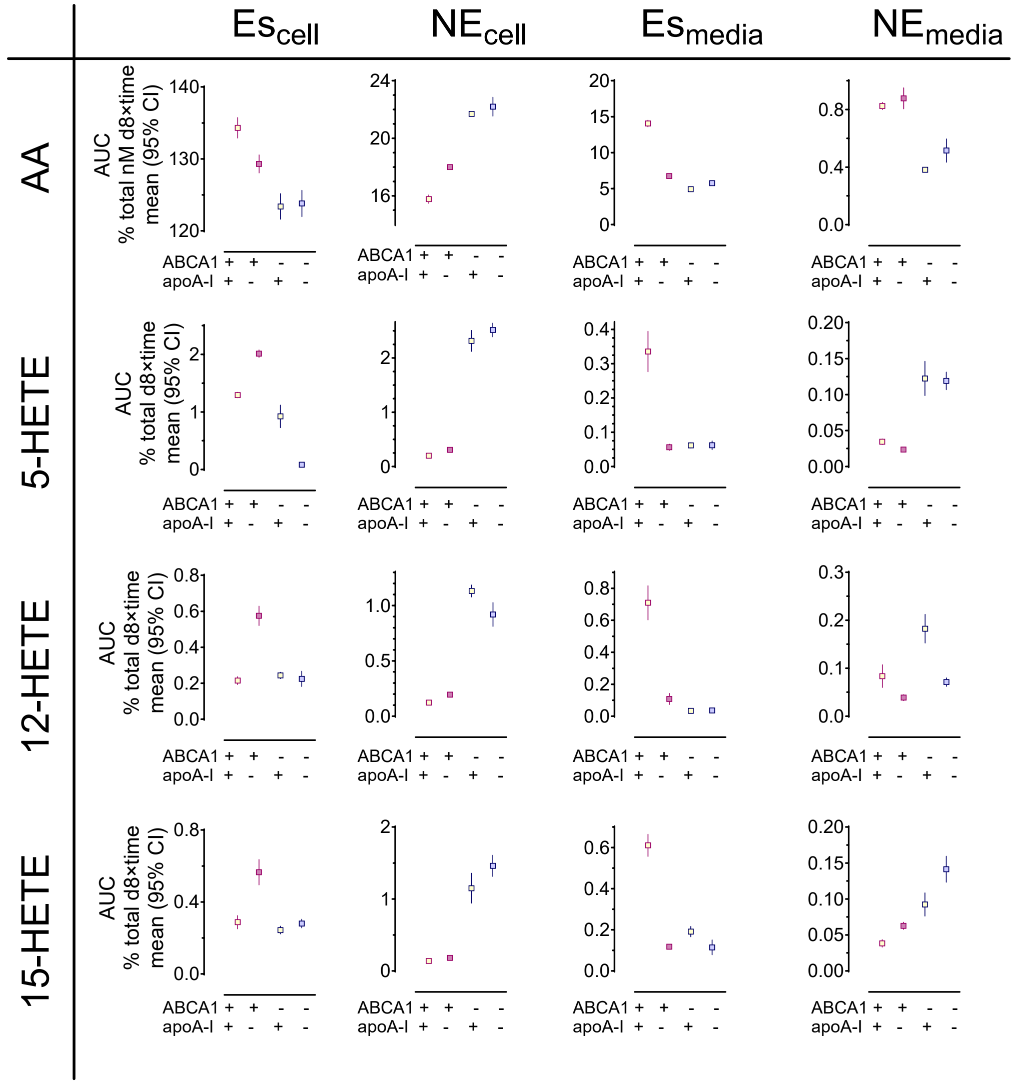

**Supplemental Figure 3: ABCA1 minimizes intracellular NE-HETE; Es-HETE in media requires combined apoA-I and ABCA1.** AUC outcomes represent cumulative exposure to the metabolite and correspond well with FTRs: 1) $NE_{cell}$-AA and $NE_{cell}$-HETE were high in the absence of ABCA1 ($1^{st}$ column), and among ABCA1 expressing cells, moderately more so in cells not exposed to apoA-I than in cells exposed. In the absence of apoA-I, ABCA1-expressing cells were able to minimize exposure $NE_{cell}$-HETE exposure by simple sequestration of excess HETEs as $Es_{cell}$-HETEs; however, when exposed to apoA-I, the cells were able to minimize both $NE_{cell}$-HETE and $Es_{cell}$-HETE. The accumulation of $NE_{media}$-HETE appeared to be weakly and inversely related to the NEcell-HETE in the case of 5- and 15-HETE, but not in the case of 12-HETE.

**Supplemental Figure 4: HETEs derived from autooxidation.**

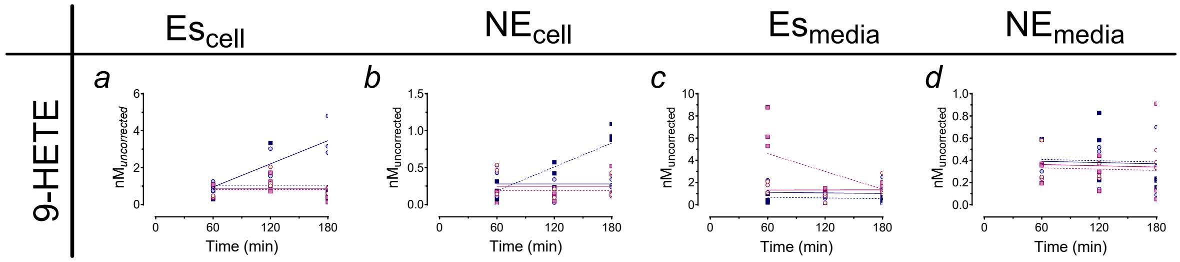

**Supplemental Figure 4: HETEs derived from autoxidation.** Unlike 5-HETE, 12-HETE, and 15-HETE which are derived from enzymatic action, oxygenation of AA at the Δ9-position results only from antioxidative activity, providing an internal control for the effects of autooxidation. Overall, 9-HETE was present at low levels compared to non-autoxidative HETEs and there was little time-dependent or condition-dependent accumulation of 9-HETE, with moderate effects (by AICc) in: 1) the $Es_{cell}$ pool where accumulation in the veh + Sil condition was moderately >0; 2) in the $NE_{cell}$ pool where accumulation in the apoA-I + Sil condition was moderately >0; and in the $Es_{media}$ pool, where a moderate rate of disappearance of 9-HETE occurred. All of the 9-HETE concentrations were comparatively low, the changes in accumulation were moderate and the effects disappeared after adjusting for multiple testing. We concluded that apoA-I/ABCA1-dependent HETE efflux was a property of enzymatically produced HETEs and did not further explore 9-HETE transfer.

**Supplement Table 3: Baseline subject characteristics (All subjects male: N=17)**

| Characteristics (Units) | Mean (±SD) |
|---|---|
| Age (yrs) | 26 (5) |
| BMI (kg/m$^2$) | 25 (2) |
| Systolic blood pressure (mg/Hg) | 118 (10) |
| Diastolic blood pressure (mg/Hg) | 77 (6) |
| Plasma glucose (mM) | 5.1 (0.4) |
| Plasma insulin (mM) | 4.5 (0.5) |
| Total cholesterol (mg/dL) | 164 (25) |
| HDL-C (mg/dL) | 52 (11) |
| LDL-C (mg/dL) | 96 (18) |
| Plasma triglycerides (mg/dL) | 81 (39) |

**Baseline characteristics of endotoxin challenge**

Baseline characteristics of participants in endotoxin challenge. Young, healthy male participants[28] had normal weights, plasma values of glucose, insulin, triglycerides, and cholesterol. Plasma TG, HCL-C, LDL-C, and glucose were all within normal values, specifically total cholesterol was < 200 mg/dL, LDL was < 100 mg/dL, and HDL > 40 mg/dL. Subjects were 65% White, 29% Asian, and 6% Black.

**Supplemental Table 4**

| Genes | Forward Primer (5' to 3') | Reverse Primer (3' to 5') |
|---|---|---|
| ABCA1 | GTCCTCTTTCCCGCATTATCTGG | AGTTCCTGGAAGGTCTTGTTCAC |
| GAPDH | AGCTTGTCATCAACGGGAAG | TTTGATGTTAGTGGGGTCTCG |

### Supplemental Table 5: Mass Transitions for MRM Analysis of Oxylipins

| Compound | Parent ion | Product ion | Dwell Time (sec) | Retention Time (mins) | Cone (V) | CE (eV) | LOD (nM) | LOQ (nM) | Internal Standard |
|---|---|---|---|---|---|---|---|---|---|
| 9-HODE d4 | 299.1 | 172.1 | 0.02 | 7.04 | 34 | 16 | 0.33 | 0.99 | - |
| 13-HODE | 295.1 | 195.1 | 0.02 | 6.99 | 34 | 14 | 0.10 | 0.31 | 9-HODE d4 |
| AA | 303.1 | 205.1 | 0.08 | 9.90 | 42 | 24 | 0.08 | 0.25 | |
| AA-d8 | 311.1 | 212.2 | 0.08 | 9.85 | 42 | 24 | 0.12 | 0.36 | |
| 12-HETE | 319.0 | 179.1 | 0.02 | 7.43 | 32 | 14 | 0.25 | 0.79 | 9-HODE d4 |
| 5-HETE | 319.1 | 114.9 | 0.08 | 7.64 | 32 | 14 | 0.29 | 0.88 | 9-HODE d4 |
| 9-HETE | 319.1 | 123.0 | 0.02 | 7.54 | 32 | 14 | 0.22 | 0.67 | 9-HODE d4 |
| 15-HETE | 319.1 | 219.0 | 0.02 | 7.17 | 32 | 14 | 0.26 | 0.79 | 9-HODE d4 |
| 5-HETE-d8 | 327.2 | 114.9 | 0.02 | 7.59 | 35 | 14 | 0.33 | 1.01 | 9-HODE d4 |
| 12-HETE-d8 | 327.2 | 184.1 | 0.02 | 7.38 | 35 | 14 | 0.31 | 0.95 | 9-HODE d4 |
| 15-HETE-d8 | 327.2 | 219.0 | 0.02 | 7.12 | 35 | 14 | 0.29 | 0.89 | 9-HODE d4 |
| CUDA | 340.2 | 214.1 | 0.05 | 5.82 | 34 | 12 | - | - | - |

**Supplemental Figure 6: Detection of HETEs and *d8*-HETEs**

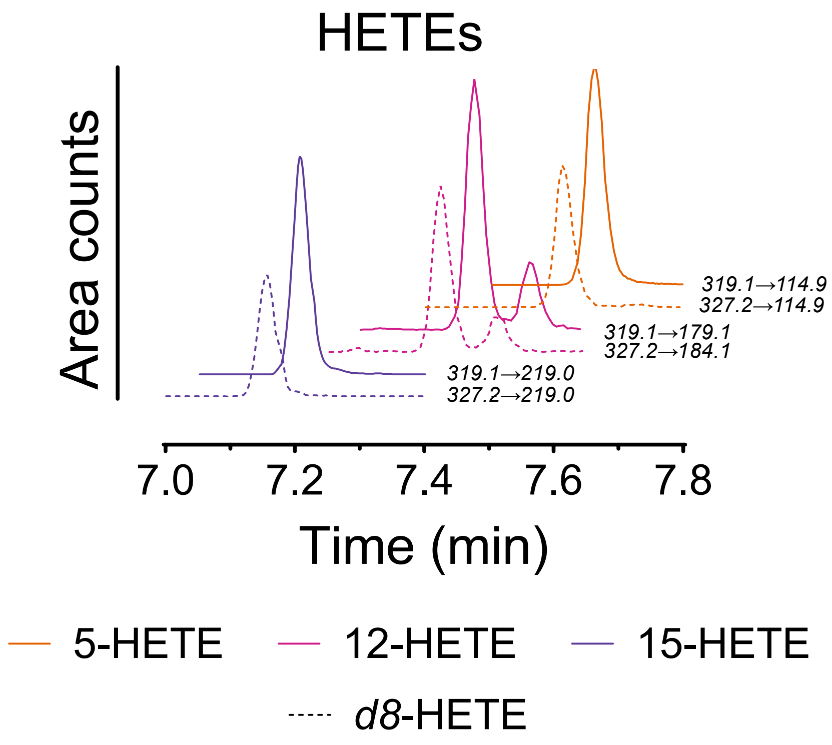

**Supplemental Figure 6 – Representative HETE traces:** Representative chromatogram traces for 5-, 12-, and 15-HETE are shown with mass transition monitored for each with parent tracee (solid) and d8 tracer (hatched). Elution of *d8*-HETEs immediately preceded the parent tracee. In all cases the response over the reported range was linear and quantitation was based on calibration curves constructed from the chromatograms.
